# Microbial Metabolite Trimethylamine N-Oxide Promotes *Campylobacter jejuni* Infection by Escalating Intestinal Inflammation, Epithelial Damage, and Barrier Disruption

**DOI:** 10.1101/2024.04.10.588895

**Authors:** Caitlyn B. Smith, Angela Gao, Paloma Bravo, Ashfaqul Alam

## Abstract

The interactions between *Campylobacter jejuni*, a critical foodborne cause of gastroenteritis, and the intestinal microbiota during infection are not completely understood. The crosstalk between *C. jejuni* and its host is impacted by the gut microbiota through mechanisms of competitive exclusion, microbial metabolites, or immune response. To investigate the role of gut microbiota on *C. jejuni* pathogenesis, we examined campylobacteriosis in the IL10KO mouse model, which was characterized by an increase in the relative abundance of intestinal proteobacteria, *E. coli*, and inflammatory cytokines during *C. jejuni* infection. We also found a significantly increased abundance of microbial metabolite Trimethylamine N-Oxide (TMAO) in the colonic lumens of IL10KO mice. We further investigated the effects of TMAO on *C. jejuni* pathogenesis. We determined that *C. jejuni* senses TMAO as a chemoattractant and the administration of TMAO promotes *C. jejuni* invasion into Caco-2 monolayers. TMAO also increased the transmigration of *C. jejuni* across polarized monolayers of Caco-2 cells, decreased TEER, and increased *C. jejuni*-mediated intestinal barrier damage. Interestingly, TMAO treatment and presence during *C. jejuni* infection of Caco-2 cells synergistically caused an increased inflammatory cytokine expression, specifically IL-1β and IL-8. These results establish that *C. jejuni* utilizes microbial metabolite TMAO for increased virulence during infection.

## INTRODUCTION

*Campylobacter jejuni* is one of the leading infectious causes of human foodborne gastroenteritis worldwide in both developed and developing nations^1^. It causes 100-150 million cases annually worldwide^2^. According to the CDC, there are about 1.5 million cases of *Campylobacter* infection each year resulting in an economic cost between $1.3 to $6.8 billion annually in the USA. *C. jejuni* has rapidly developed multiple mechanisms of antibiotic resistance all over the world^3,4^. Hence, the CDC designated the drug-resistant *Campylobacter* strains as a “serious antibiotic resistance threat.” Therefore, *C.jejuni* warrants deeper investigation into the mechanisms of its pathogenesis, which may allow the development of potential therapeutic agents against it.

As a microaerobic human enteric pathogen, *C. jejuni* is a Gram-negative, proteolytic, spiral bacterium belonging to the Campylobacterota phylum^1^. During infections in humans, *C. jejuni* passes through the stomach and small intestine to colonize within the distal ileum and ascending colon, and also traverse the space from the anaerobic intestinal lumen to the microaerobic crypts of the epithelium, which is lined with viscous mucins^3,5^. *C. jejuni* specifically penetrates and colonizes the mucin layer, eventually establishing adherence and invasion of the gut epithelium layer^3,6^, which is a dynamic barrier segregating the luminal contents and gut microbiota from the underlying mucosal tissue compartments^7,8^. Epithelial barrier integrity is regulated by intercellular junctions, including the tight junction (TJ), adherens junction, and desmosomes. Disruption of epithelial TJs may result in the translocation of luminal contents, bacteria, and bacterial products into the mucosal compartments, leading to intestinal and extraintestinal inflammation^9,10^.

To establish infection and cause gastroenteritis in humans, *C. jejuni* employs a repertoire of factors, which include flagella for chemotaxis and colonization of the mucus layer, several adhesins and other proteins for adherence and invasion of cells, as well as a single exotoxin called cytolethal distending toxin (CDT)^4,11^. In addition, *C. jejuni* disrupts epithelial paracellular junctions in several ways, including secreting serine protease HtrA, switching claudin isoforms, or redistributing TJ protein cellular location^12^. *C. jejuni* infection also causes intestinal epithelial barrier dysfunction by decreasing claudin 4 level or increasing pore-forming claudin 2 expression and by focal redistribution of occludin and TJ proteins^13^. This damage to the intestinal epithelium causes infected individuals to have diarrhea containing blood and leukocytes, abdominal cramping, vomiting, and fever. In developing countries, campylobacteriosis in children under the age of 2 years is highly prevalent, sometimes resulting in death^3^. Though *C. jejuni* infections are self-limiting, this disease can cause many secondary diseases, such as acute autoimmune diseases, including arthritis and the neuropathies Guillain-Barré syndrome and Miller Fisher Syndrome. Furthermore, the inflammation caused by *C. jejuni* has been implicated with irritable bowel syndrome and inflammatory bowel disease in some patients, an inflammatory condition that has been linked to cancer^14^.

Human *C. jejuni* infections are acquired from contaminated reservoirs or food, such as water, poultry, swine, and bovine sources^3^. Avian guts are often colonized by *C. jejuni* as a commensal bacterium at a high level (10^6^ to 10^8^ cfu/g). Interestingly, this zoonotic bacterial pathogen colonizes the gastrointestinal tract of wild and domestic animals asymptotically without causing significant macroscopical and microscopical lesions. In contrast, infection with a dose as low as 100 CFU may result in diarrhea in humans. However, it remains poorly understood how *C. jejuni* successfully establishes commensalism in many animal hosts but promotes diarrheal disease in the human population^14^. Furthermore, it is unclear why *C. jejuni* infections lead to predominantly inflammatory, bloody diarrhea, dysentery syndrome in developed countries, but asymptomatic infections or watery diarrhea in developing nations. It is hypothesized that gut microbiota and their metabolites of the different host intestines are unique, which function as the key determinants for *C. jejuni* to sense and reprogram its virulence factors to exert either commensalism, asymptomatic infections, or pathogenesis of diarrheal disease^15,16^. Interestingly, *C. jejuni* has a relatively small genome compared to its worldwide incidence and pathogenicity, of about 1.7 megabase pairs, translating to about ∼1600 genes, making the pathogen a major scavenger^17,18^. Although *C. jejuni* has a small genome and, thus, a reduced metabolic capacity, the pathogenic mechanisms of *C. jejuni* is not completely understood. Unlike other enteric pathogens with more than 4,500 genes, we posit that a minimal genome allows *C. jejuni* to vigorously acquire small molecules from the environment for its growth, expansion, and successful colonization in the host intestine. Thus, it is plausible that *C. jejuni* obtains and perceives the abundance of specific commensal microbial metabolites to rewire its metabolic pathways and virulence gene expression networks according to the types of the host it colonizes and the biogeographic location in that gastrointestinal tract.

Recent research studies suggest that the intestinal microbiota and microbial metabolites impact enteropathogenesis^19^, and this mechanism of action differs based on the pathogen. For example, the Lactobacilli genus conveys colonization resistance to *Shigella* species by increasing the anti-inflammatory state, which blocks the pathogen from adhering to the epithelial cells ^20,21^. Similarly, Commensal *Enterobacteriaceae* species convey colonization resistance against *Salmonella enterica* serovar enteritidis by competing for O_2_ in the environment^22^. As mentioned previously, *C. jejuni* infections are mostly localized at the distal ileum and colon in the human host, which are the physiological location of the bulk mass of the gut microbiota. As an enteric pathogen, *C. jejuni* has a unique relationship with the intestinal microbiota. The C57Bl/6 mouse model, when maintained in standard laboratory (specific pathogen free) conditions, displays colonization resistance to *C. jejuni*^23^, a protection against colonization of an organism provided by the intestinal microbiota via systems such as the production of antimicrobial molecules, the activation of immune systems, the competition for physical space, and the use of microbial-produced metabolites^24^. Furthermore, this protection against *C. jejuni* in C57Bl/6 mice is abrogated during intestinal dysbiosis, and this dysbiosis can be inflicted simply with dietary modifications or antibiotics^23^. Presently, we do not completely understand the interactions among *C. jejuni,* gut microbiota, and microbial metabolites. It has been reported previously that the primary and microbially modified secondary bile acids in the small intestine cause differential protein loading in the outer membrane vesicles of *C. jejuni,* which may result in increased epithelial damage^25,26^. Furthermore, studies comparing *C. jejuni* infectible and resistant mouse models demonstrate that *C. jejuni* infection is associated with increased Enterobacteria, Enterococci, *Bacteroides/Prevotella* spp., particularly increased *Escherichia coli*, and decreased Lactobacilli and mouse intestinal Bacteroides^27^. Additionally, it was demonstrated that the intestine of the mouse model, which is resistant to *C. jejuni* infection, contained high levels of antimicrobial fatty acids as well as a low level of amino acids commonly utilized by *C. jejuni* for growth and expansion. Furthermore, the gut metabolite L-lactate plays a vital role as a growth substrate for *C. jejuni* during acute infection^6^. Recent studies have begun deciphering the importance of the gut microbiota and its metabolites during *C. jejuni* infection; however, we still do not completely comprehend their functional mechanisms.

*C. jejuni* utilizes chemotaxis systems to navigate and to mediate directional motility towards the mucus and intestinal crypts for host infection. The chemotaxis systems of *C. jejuni* are similar to the canonical two-component chemotaxis systems: methyl-accepting chemotaxis proteins (MCP). It is usually called transducer like proteins (Tlp) in *C. jejuni*, which help the bacterium to sense extracellular molecules (chemoeffectors/ligands)^28,29^. Whether Tlp senses a chemoattractant or chemorepellent, this signal gets relayed to the histidine kinase, CheA, which is held to the MCP by *Campylobacter-*specific scaffolding proteins called CheW and CheV. Subsequently, CheA acts upon a Response Regulatory (RR) protein CheY. Phosphorylated CheY interacts with the flagellar motor, which changes the direction of rotation between clockwise and counterclockwise. This alteration enables the organism to move towards an attractant or away from a repellant. *Campylobacter* species encodes for upwards of 25 different MCPs, which also includes systems for energy taxis and aero taxis. Nearly all the MCPs have multiple different chemoeffectors. As with other pathogens, this chemotaxis is essential for pathogenesis in the host. Single deletions of multiple different MCPs in *C. jejuni* demonstrated that this chemotaxis is necessary for exiting the intestinal lumen and swimming toward the intestinal mucosa. In addition, these *C. jejuni* mutants with defective Tlps are incapable of invading epithelial cells *in vitro* and also demonstrate reduced colonization of mouse colonic mucosal tissues^30–32^.

In humans and mice, intestinal dysbiosis can cause the expansion of the Proteobacteria phylum and increase the production of biogenic amines^33–35^. These biogenic amines are either derived from protein-rich diet or arise as the byproducts of the microbial degradation of proteins and amino acids. Diet-derived biogenic amines are mostly absorbed in the small intestine, and thus, the colonic biogenic amines are produced mostly by the microbiota^36^. One key microbial metabolite that becomes highly enriched during gut dysbiosis is trimethylamine *N*-oxide (TMAO). The dysbiotic gut microbiota, especially proteobacteria, including *E. coli*, catabolize choline, carnitine, and betaine into the precursor TMA^37–39^, which is converted to TMAO by flavin monooxygenase 3 (FMO3) in the host liver^38,40^. Human studies demonstrated that the serum levels of TMAO is directly correlated to the increased intestinal levels of the Proteobacteria phylum and Bacteroides phylum^41,42^. Not only are germ free mice devoid of TMAO within their serum^43^, but the therapeutics intervention to reduce the TMAO levels in humans mostly target the bacterial production of this biogenic amine^44^, emphasizing the role of the gut microbiota in TMAO production. TMAO has also been associated with human health and many diseases^45^, including cardiovascular diseases, cancer, and arthritis. Recently, TMAO has been shown to affect multiple other pathogens, such as *Vibrio cholerae* by increasing the anaerobic cholera toxin production and modulating *in vivo* colonization^45,46^. In addition, TMAO promotes *Helicobacter pylori* pathogenesis by synergistically increasing the inflammatory response of gastric epithelial cells^47^. Also, the colonization of uropathogenic *Escherichia coli* to the bladder epithelial cells and subsequent inflammatory responses are critically affected by TMAO^48^. These results suggest that TMAO can synergistically enhance bacterial colonization and pathogen-mediated inflammation *in vitro and in vivo*.

As a major scavenger, *C. jejuni* infects the distal ileum and colon, which is heavily enriched with microbiota and its metabolites. *C. jejuni* genome encodes a TMAO reductase, termed *torA*, which enables the pathogen to utilize TMAO as a terminal electron acceptor and produce energy in anaerobic conditions found within the colonic lumen^49^. *C. jejuni* strictly regulates its respiration systems and energy production. In the presence of TMAO, the two-component system RacSR regulates the expression of specific electron donor systems for fumarate (*mfr),* aspartate (*aspA),* C4-dicarboxylates (*dcuA),* cytosolic glutamate (*gltBD)* and γ-glutamyltranspeptidase (*ggt)*^50^. The mutant *C. jejuni* with a *torA* gene deletion has demonstrated decreased colonization in a broiler chicken model^51^. However, how TMAO influences the pathogenesis of *C. jejuni* is not completely understood.

There is a knowledge gap in our understanding of the effect of the gut microbiota and their metabolites on *C. jejuni* infection. Hence, the questions we examined in this study include: 1) Does *C. jejuni* infection in mice cause alterations in gut microbiota-associated production of specific biogenic amines in the intestinal lumen? 2) Does microbial inflammatory biogenic amine TMAO influence *C. jejuni* pathogenesis? 3) Does TMAO impact *C. jejuni* by causing TJ dysfunction and disrupting the epithelial barrier?

Thus, our main goal in this study is to understand the relationship between *C. jejuni* 81-176 and microbial metabolites during infection. Here, we determined that a *C. jejuni* infection resulted in a remarkable microbial dysbiosis in IL10KO mice, which is characterized by elevated levels of Proteobacteria, *E. coli,* and inflammatory markers. As hypothesized, *C. jejuni* infection also increased TMAO levels within the colonic lumens of IL10KO mice but not in WT mice. We further experimentally determined that TMAO treatment promoted bacterial chemotaxis, invasion, and transmigration of *C. jejuni in vitro*, as well as enhanced *C. jejuni-*mediated epithelial damage and inflammatory cytokine expression.

## RESULTS

### *C. jejuni* infection escalated gut microbiota alterations and elevated inflammatory markers in IL10KO mice

Recent research studies suggest that the intestinal microbiota and microbial metabolites impact the enteropathogenesis of bacterial pathogens. To expand our understanding of the effect of the gut microbiota and their metabolites on *C. jejuni* infection, first, we used IL10KO mice as conventionally raised C57Bl/6 mice exhibit colonization resistance to *C. jejuni* ^23,24,52^. Six-week-old WT and IL10KO mice were gavaged either 5×10^9^ CFU of *C. jejuni* strain 81-176 or 0.3 mL of 1x PBS as a vehicle control for two consecutive days. The mice were then monitored over the span of 2 weeks before termination. Using the stool samples, the microbial compositions were analyzed in both male and female mice by sequencing^7,53^ 16S rRNA gene amplicons in WT and IL10KO mice with or without *C. jejuni* infections (Fig. 1). At the phyla level, microbiota analysis revealed that *C. jejuni* infection remarkably caused dysbiosis in the intestinal microbiota (Fig. 1A). A principal coordinates plot (PCoA) also demonstrated that the microbiota associated with *C. jejuni*-infected IL10KO mice clustered distinctly from that of uninfected IL10KO mice (Fig. 1B). We also determined that there was a significant increase in the Proteobacteria phylum and relative abundance of *E. coli* in the uninfected IL10KO mice compared to the WT mice (Fig. 1A, C,D). Importantly, we found that *C. jejuni* infection remarkably expanded the relative abundance of Proteobacteria phylum (Fig. 1C) and *E. coli* species (Fig. 1D) in the IL10KO mice but not in the WT mice, reciprocating past literature on the correlation between increased intestinal *E. coli* and *C. jejuni* susceptibility^54–56^. Additionally, *C. jejuni* infection reduced the relative abundances of Actinobacteria, Verrucomicrobia and Firmicutes phyla in the IL10KO mice (Fig. 1A). We also recovered significantly elevated levels of *C. jejuni* (CFU/g of stool) in the IL10KO mice compared to the WT mice. We next evaluated the levels of inflammation markers lipocalin-2 (LCN-2) in the stool and tumor necrosis factor α (TNF-α) in the colonic tissues. LCN-2, also called neutrophil gelatinase-associated lipocalin (NGAL), functions as an iron siderophore to outcompete pathogens and is a secreted biomarker of inflammation and infection and found in association with *C. jejuni* infections in stool ^57–59^. TNF-α is a proinflammatory cytokine with major roles in gut health but also in initiating the innate immune response to *C. jejuni* infection ^60,61^. We found that both LCN-2 (Fig. 1F) and TNF-α (Fig. 1G) accumulation is significantly exacerbated in the IL10KO mice during *C. jejuni* infection as compared to the uninfected ILKO mice or infected WT mice.

**Figure 1.**
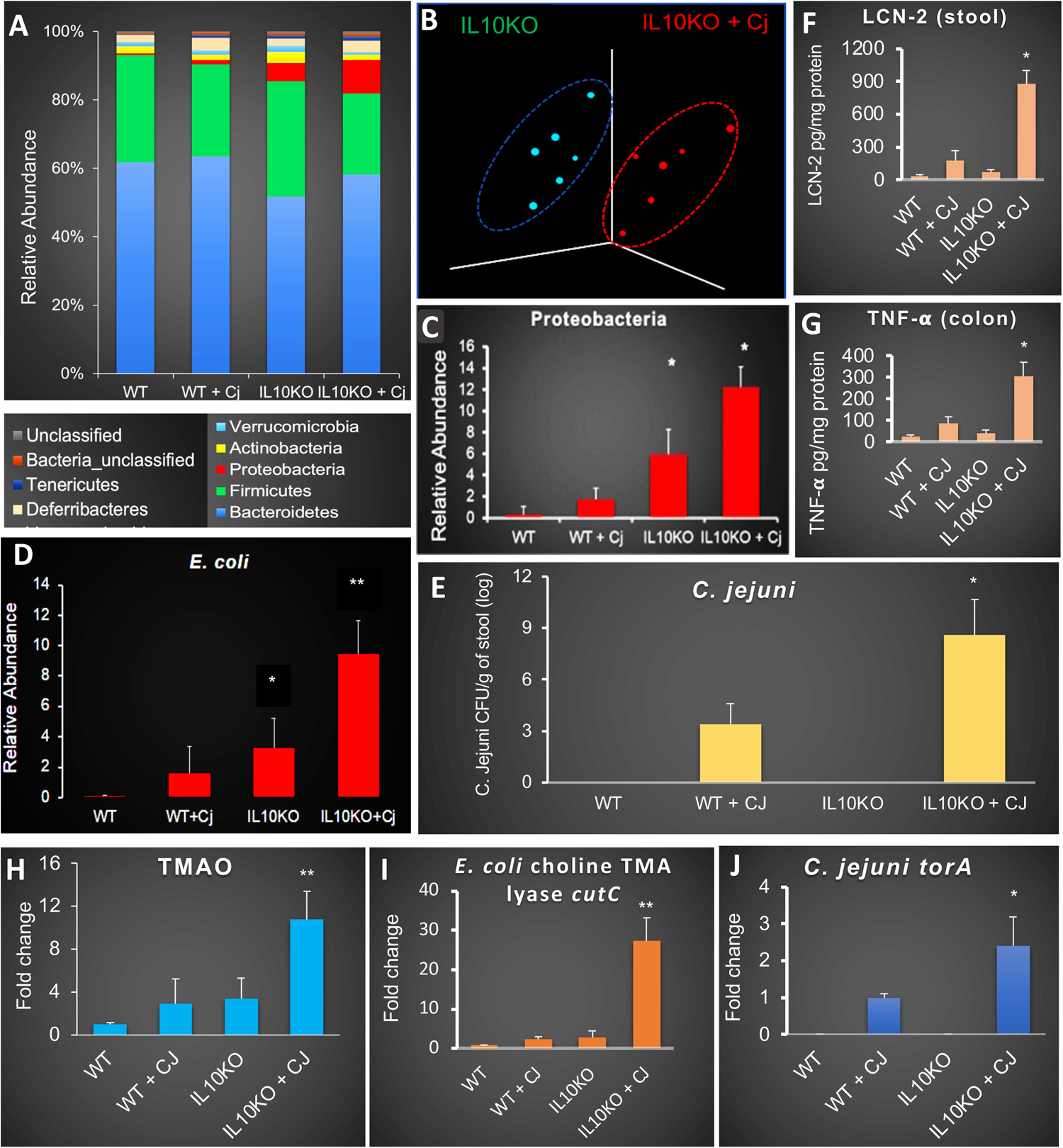
*C. jejuni* infection induces alterations of gut microbiota and inflammatory markers in IL10KO mice. (A) Microbial community analysis and mean relative abundances of bacterial phyla. Mice were gavaged with 5×10^9^ *C. jejuni* 81-176 CFU/mouse for 2 consecutive days. 14 dpi, stool, luminal contents, and colons were harvested. Bacterial 16s rRNA profiling was done by high-throughput sequencing (HTS). **CJ** denotes *C. jejuni* (B) PCoA plot of microbiota community structure in uninfected IL10KO mice and IL10KO mice infected with *C. jejuni*. Microbial community analysis and mean relative abundances of (C) Proteobacteria phylum and (D) *E. coli* in the mice, determined by HTS of 16S rRNA. (E) CFU enumeration of *C. jejuni* in the stool of mice. (F) LCN-2 in stool and (G) TNF-α in colonic tissues were measured via ELISA. (H) Fold change of TMAO determined by LC-MS of luminal content. (I-J) qPCR analysis of the *E. coli* choline TMA lyase *cutC* gene (I) and *C. jejuni* Tor reductase *torA* gene (J). Data are presented as the mean ± SEM of two independent biological replicates, (n=8). Statistical significance was assessed via one-way ANOVA followed by Fisher’s LSD post hoc test. * p <0.05 ** p<0.01

### *C. jejuni* infection elevated bacterial metabolite TMAO in the luminal content in IL10KO

Gut microbiota and its metabolites profoundly impact the pathogenesis of enteric bacteria, yet the exact mechanisms by which gut commensals’ metabolic alterations influence *C. jejuni* infection and virulence remain incompletely understood. As a microaerobic enteric pathogen, *C. jejuni* requires terminal electron acceptors in the intestinal microenvironment. One of the electron acceptors that commensal bacteria contribute to the intestinal microenvironment is TMAO. Gut microbiota, especially proteobacteria, including *E. coli*, catabolize choline into precursor TMA^37–39^, which is converted to TMAO by flavin monooxygenase 3 (FMO3) in the liver^38^. TMA/TMAO is often a source of energy for other microorganisms ^39^. Interestingly, *E. coli* has been shown to be associated with elevated choline catabolism and TMAO enrichment^61^. TMAO is an inflammatory metabolite that can be used by *C. jejuni* TMAO reductase, TorA^49^. Our microbial community survey uncovered that *C. jejuni* infection significantly increased the relative abundance of *E. coli* in the IL10KO mice (Fig. 1D). Hence, we investigated whether *C. jejuni* infection promotes TMAO abundances by conducting LC-MS/MS analysis of the luminal contents collected from the mice. We found higher levels (not significant) of TMAO in both IL10KO, and WT mice infected with *C. jejuni* than in uninfected WT mice. Remarkably, *C. jejuni* infection significantly elevated the TMAO level in the IL10KO mice (Fig. 1H). Interestingly, the abundance of mRNA of *E. coli*’s choline TMA lyase *cutC* gene is also dramatically increased in IL10KO mice infected with *C. jejuni* but not in other groups of mice. BLAST search and *C. jejuni* genome analysis confirmed that *C. jejuni* 81-176 does not encode a homologous gene to *E. coli*’s *cutC* gene (data not shown). Furthermore, the *C. jejuni* mRNA encoding the TorA enzyme for TMAO reduction is also significantly elevated in *C. jejuni*-infected IL10KO mice, indicating an important role of elevated levels of *E. coli* and TMAO metabolites during *C. jejuni* infection of IL10KO mice.

### *C. jejuni* senses TMAO and precursors to TMAO as chemoattractant

Chemotaxis enhances the ability of *C. jejuni* to navigate its host environment to obtain and metabolize substrates, escape noxious compounds/environment, adhere to colonocytes, and transmigrate into mucosal tissues and beyond, thereby promoting successful colonization and infection^30–32^. *C. jejuni* is a microaerobic bacteria and utilizes TMAO as an alternative electron acceptor under oxygen-limiting conditions^51^, predominantly found within the colonic lumen of the hosts^50^. However, it is not known whether *C. jejuni* senses TMAO as a chemoeffector. To evaluate the effect of TMAO on *C. jejuni’s* chemotaxis, we conducted a chemoattractant-based semisolid (0.3% agar) stab-agar assay^62^. In addition to the vehicle control and serine, a known chemoattractant for *C. jejuni* ^31,63,64^, we examined TMAO, as well as other precursor molecules of TMA, including L-carnitine and betaine, found in red meat and proteinaceous leafy greens, respectively^37,65^. After 24 hours of incubation under microaerobic conditions, the zones were measured according to Mo et al.^66,67^. We found that, in comparison with vehicle control, TMAO and its precursor molecules, including L-carnitine and betaine, promoted chemotaxis statistically similar to serine, a known strong chemoattractant^68^ for *C. jejuni* (Fig. 2A-B). Here, we show that *C. jejuni* senses TMAO as a chemoattractant in microaerobic conditions and can promote *C. jejuni* chemotaxis.

**Figure 2.**
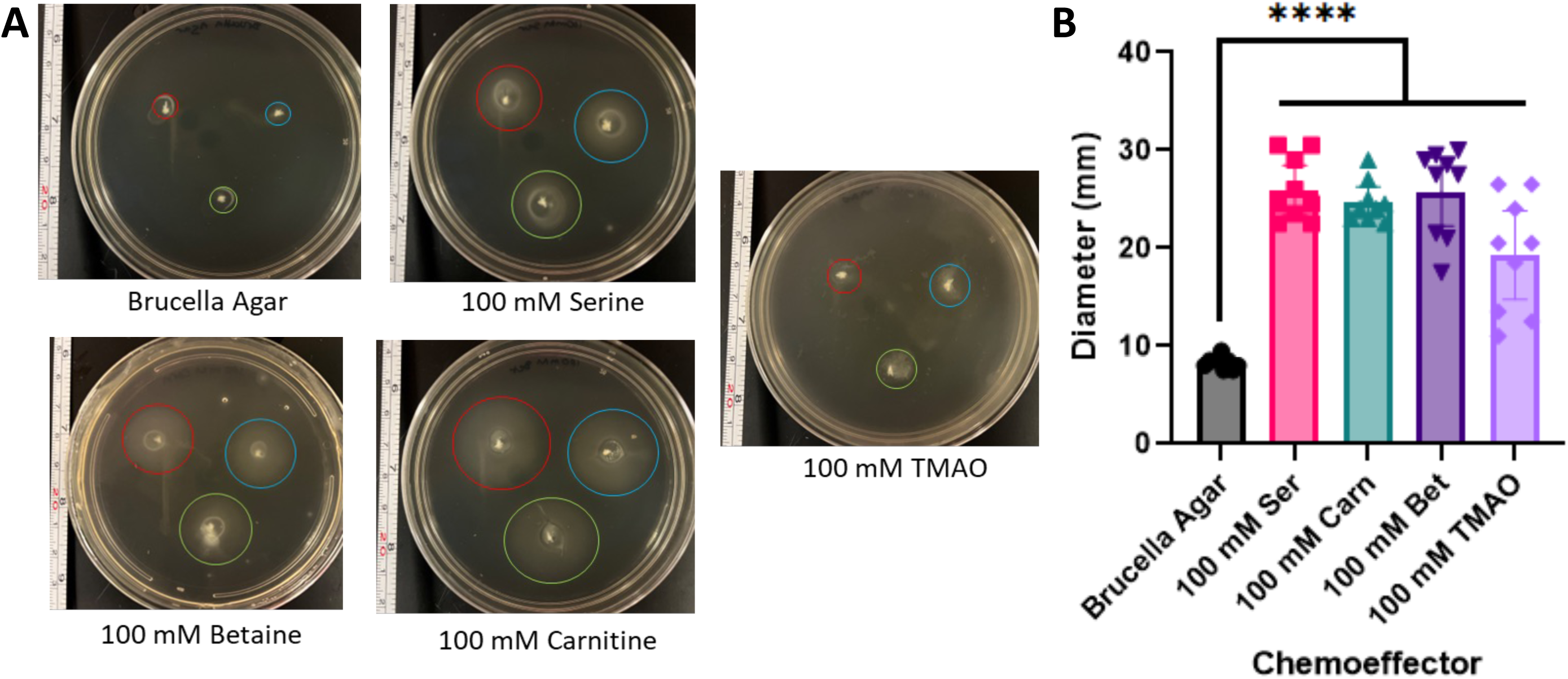
Microbial Metabolite TMAO is a chemoattractant for *C. jejuni*. (A) Representative images, and (B) results from 3 independent stab agar chemotaxis assays of TMAO and other precursor molecules compared to controls. Data are presented as the mean ± SEM of three independent biological replicates. Statistical significance was assessed via (B) One-Way ANOVA followed by Dunnett’s post hoc test. **** p<0.0001

### Bacterial biogenic amine TMAO increases *C. jejuni* invasion *in vitro*

After reaching the colon, *C. jejuni* next colonizes & invades the mucosa and enterocytes as was demonstrated in biopsies of infected patients and using *in vitro* infection assays^69^. After determining that TMAO influences the chemotaxis of *C. jejuni*, we next examined whether this metabolite could also influence its virulence mechanisms. To test this, we utilized Caco-2 cells, a human colorectal adenocarcinoma cell line that mimics the colonic epithelial layer^69–71^. We cultured Caco-2 cell monolayers to reach confluency, and monolayers were then treated with varied concentrations of TMAO for a total of 48 hours. We investigated the invasion of *C. jejuni* in three different conditions: 1) invasion assay in the presence of TMAO to observe the effects of TMAO on *C. jejuni*, 2) invasion assay in the absence of TMAO by replacing the treatment media with conditioned media to observe the effects of TMAO on the epithelium, and 3) invasion assay in the presence of TMAO under microaerobic conditions, as TMAO is utilized by *C. jejuni* as an alternative electron acceptor only when oxygen is not as readily available^50,51,72^. After the *C. jejuni* infection (MOI of 200)^16^ in the presence of TMAO for 48 hours, the Caco2 cells were treated with gentamicin as previously described^73,74^. We found that TMAO (Fig. S1A in the supplemental material) did not increase *C. jejuni* invasion in comparison to the control, suggesting that TMAO treatment is not effective in a standard aerobic culture condition. After 48 hours of TMAO treatment, next we washed the TMAO from the media and allowed the *C. jejuni* infection to continue in the absence of TMAO (Fig. S1B). We did not find any significant differences in the invasion of *C. jejuni*. When *C. jejuni* was allowed to invade in the presence of TMAO under microaerobic conditions (Fig. 3A), we found that there was significantly more *C. jejuni* recovered from the monolayer at 800 μM TMAO. Furthermore, we determined a dose-dependent increase in *C. jejuni* invasion at higher TMAO concentrations, including 1 mM, 1.25 mM, and 1.5 mM TMAO. The growth of *C. jejuni* was previously reported to be assessed in the presence of 20 −50 mM TMAO; we postulate that the lower concentrations of TMAO used in the current experiments closely reflect physiologic conditions. We further assessed if this invasion required CDT, a major virulence factor of *C. jejuni.* CDT is a toxin that causes cell cycle arrest at the G2/M phase and subsequent apoptosis of host cells ^75,76^. We conducted the invasion assay in the presence of TMAO under microaerobic conditions with a CDT deletion mutant (Δ*cdtABC*) of *C. jejuni* (Fig. 3B). We found a similar level of invasion of Δ*cdtABC C. jejuni* mutant into the Caco-2 monolayer, suggesting that this TMAO-induced *C. jejuni* invasion is not dependent on *C. jejuni*’s production of CDT.

**Figure 3.**
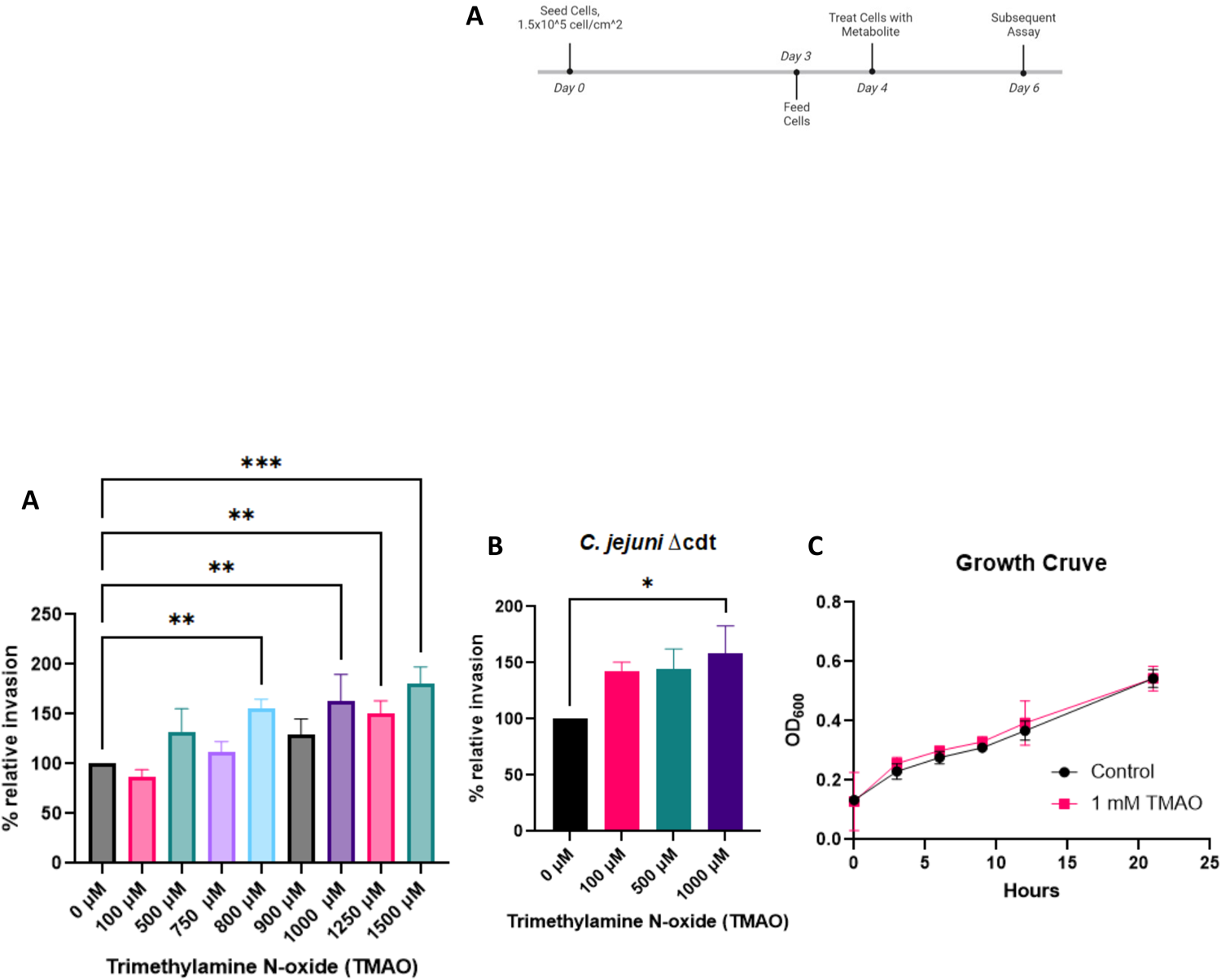
Microbial Metabolite TMAO increases the *in vitro* invasion of *C. jejuni.* Results of microaerobic invasion assay of *C. jejuni* in the presence of TMAO for (A) wild type strain 81-176 and (B) Δ*cdt* mutant deletion strain of 81-176. (C) Growth Curve of *C. jejuni* 81-176 in the presence of TMAO under microaerobic conditions. Data are presented as the mean ± SEM of three independent biological replicates. Statistical significance was assessed via (A and B) One-Way ANOVA followed by Fisher’s LSD post hoc test, and (C) Two-way ANOVA followed by Sidak’s post hoc. * p<0.05, **p>0.01, ***p>0.001

As the increase in invasion may be affected by the growth of *C. jejuni*, we also examined the growth rate of *C. jejuni* in the presence of 1 mM TMAO. The results indicated that there was no significant change in the growth rate based on the presence of TMAO under microaerobic conditions (Fig. 3C), which is in agreement with previous literature^49^. These results indicate that TMAO does not influence *C. jejuni* growth under these conditions yet enhances *C. jejuni* to invade Caco-2 cells *in vitro*.

### TMAO disrupts tight junction proteins and promotes *C. jejuni* invasion in monolayers of Caco-2 cells

Proteins that comprise the tight junctional (TJ) complexes between cells are required to convey intestinal epithelial barrier integrity, structure, and intestinal permeability^77^. The epithelial TJ is the apical-most intercellular junction and serves as a dynamic gatekeeper for the paracellular pathway. *C. jejuni* invades intestinal epithelial cells during infection^1,74,78^, but also transmigrate across the intestinal epithelium, infect basolaterally and disseminate to other organs^79–82^. *C. jejuni* is reported to use paracellular transmigration to translocate into the host mucosal tissue and invade gut epithelial cells in basolateral fibronectin & integrin-dependent manner^83,84^. For this purpose, *C. jejuni* has been demonstrated to damage epithelial barrier integrity in several ways, including secreting serine protease, switching claudin isoforms, redistributing TJ protein cellular location, or modulating TJ expression^13^. During the invasion, *C. jejuni* cleaves tight junction stabilizing protein occludin^73,84^, damaging tight junctional complexes and causing cytoskeletal rearrangement for invasion into cells^74,78,85,86^. Furthermore, *C. jejuni* infection also causes intestinal epithelial barrier disruption by decreasing claudin 4 level or increasing pore-forming claudin 2 expression and by focal redistribution of occludin and TJ proteins^74,78,85,86^.

For these experiments, we wanted not only to recapitulate our invasion assay data using an immunofluorescence and microscope-based technique but also to determine if TMAO impacts the functions of the tight junction proteins within the Caco-2 cells by causing focal redistribution from the cellular junctions that may enable *C. jejuni* invasion. To investigate this, we conducted immunofluorescence (IF) staining of Caco-2 cells treated with 1 mM TMAO for 48 hours, followed by treatment with *C. jejuni* or vehicle control (Fig. 4, Fig. 5, Fig. S2) for 3 hours. Finally, cells were treated with gentamycin. First, we determined an increase in *C. jejuni* invasion in the presence of 1 mM TMAO under microaerobic conditions (Fig. 4D), which supported our findings of the gentamycin-invasion assays (Fig. 3).

**Figure 4.**
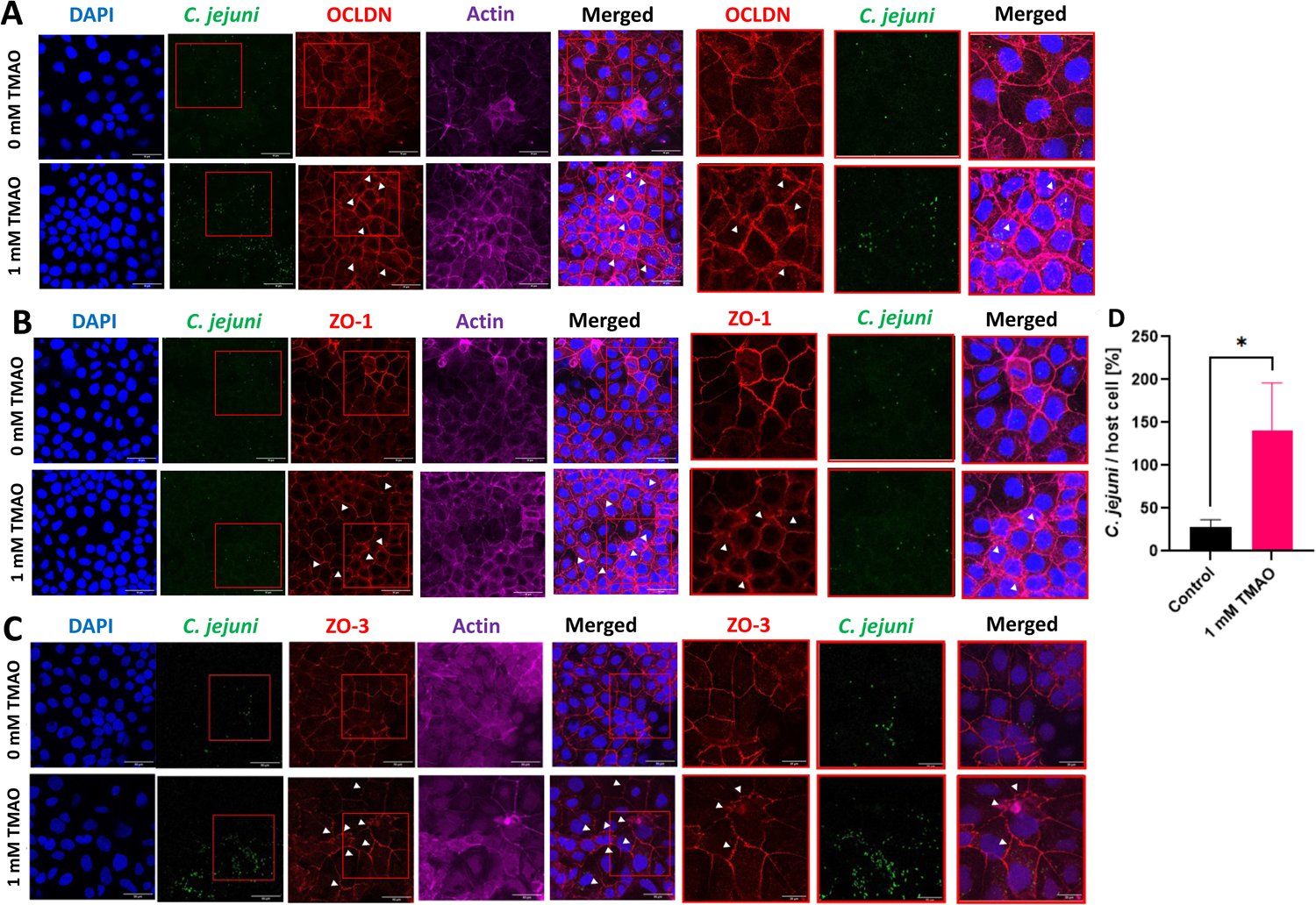
Microbial Metabolite TMAO increases the *in vitro* invasion of *C. jejuni,* visualized by immunofluorescence staining. (A-C) Representative images from fixed cells and immunofluorescence staining of the nucleus (DAPI), tight junctional complex protein (red), Actin (purple), and *C. jejuni* (green). IF staining was conducted under 2 different experimental conditions: *C. jejuni* infected untreated control and *C. jejuni* infected TMAO-treated cells. Cells were treated with TMAO for 48 hours and the 3-hour infection with *C. jejuni* was conducted under microaerobic conditions. White arrows designate sites of dislocation of tight junctional proteins from the cell-to-cell junctions, denoting loss of function. White arrows designate sites of membrane loss, ruffling or punctation observed with the representative tight junctional protein staining. Tight junctional complex proteins tested: (A) occludin (OCLDN), (B) zonula occludens-1 (ZO-1), and (C) zonula occludens-3 (ZO-3). (D) Quantification intracellular *C. jejuni* 81-176. Data are presented as the mean ± SEM of three independent biological replicates. Statistical significance was assessed via (G) Student’s t-test. * p<0.05

**Figure 5.**
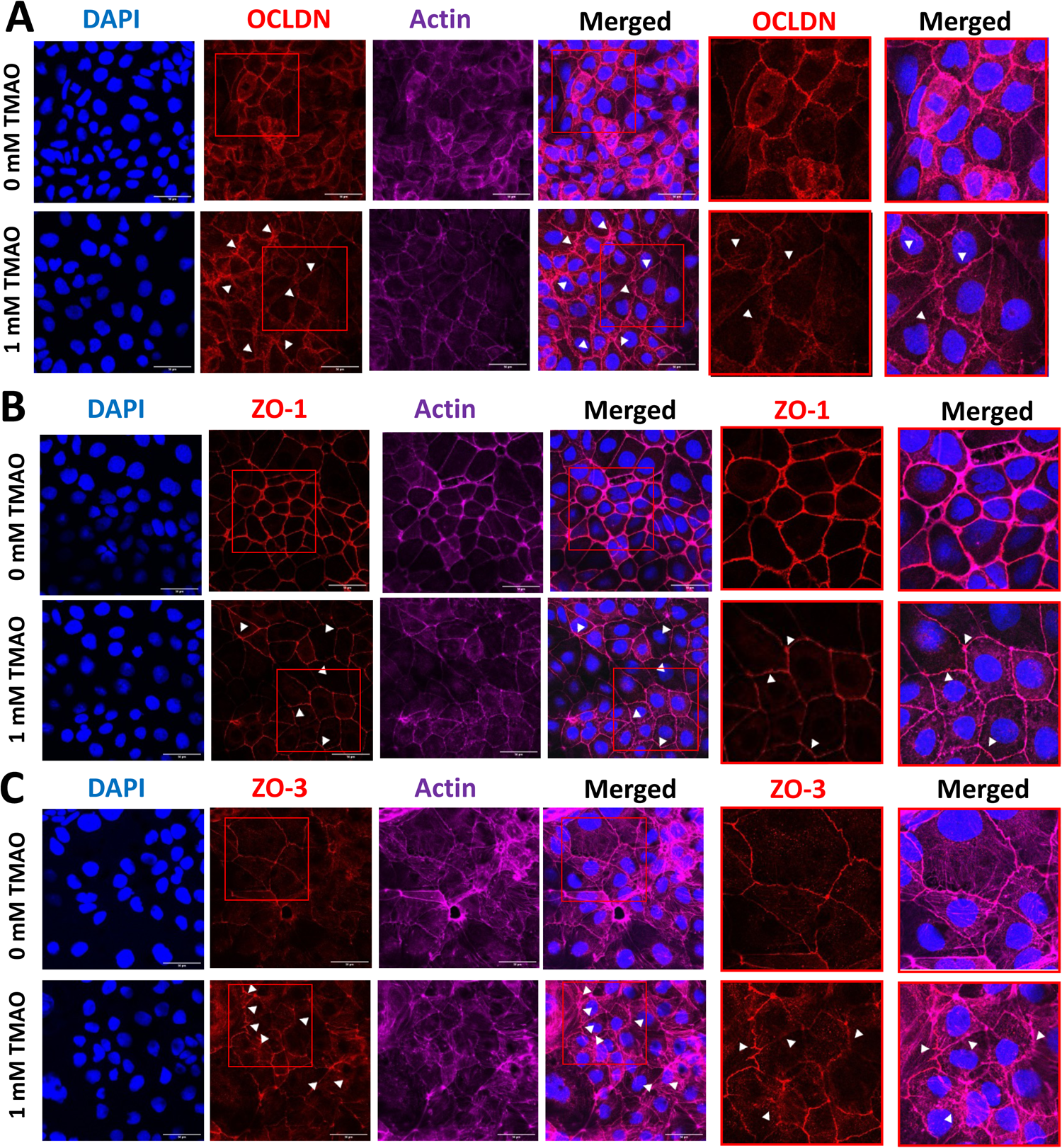
TMAO affects tight junctional complex protein function in Caco-2 cells. (A-C) Representative images from fixed cells and immunofluorescence staining of the nucleus (DAPI), tight junctional complex protein (red), and Actin (purple). IF staining was conducted under two different experimental conditions: no-treatment control, TMAO-treated cells. Cells were treated with TMAO for 48 hours including a terminal 3-hour microaerobic incubation but without *C. jejuni* infection. White arrows designate sites of dislocation of tight junctional proteins from the cell-to-cell junctions, denoting loss of function. White arrows designate sites of membrane loss, ruffling or punctation observed with the representative tight junctional protein staining. Tight junctional complex proteins tested: (A) occludin (OCLDN), (B) zonula occludens-1 (ZO-1), and (C) zonula occludens-3 (ZO-3). Data are presented as the mean ± SEM of three independent biological replicates.

Next, we examined an array of tight junctional complex proteins, including the ‘tight claudin’ claudin-4 (CLDN4), the tetraspan junctional stabilizing protein occludin (OCLDN), and intracellular scaffolding proteins zonula occludens 1 (ZO-1), zonula occludens 2 (ZO-2), and zonula occludens 3 (ZO-3). There was no significant effect on proteins claudin-4 (and ZO-2 (Fig. S2A-D) in response to the TMAO treatment. However, claudin-4 protein was dramatically diminished in response to the *C. jejuni* infection, with or without TMAO treatment (Fig. S2C). When we examined occludin, ZO-1 and ZO-3 (Fig. 4A-C), we found that in the presence of TMAO, *C. jejuni* infection caused ruffling of the cell-to-cell junctions between neighboring cells, punctation within the junctional complexes, and heightened redistribution to the cytoplasm. TJ ruffling frequently correlates with increased paracellular permeability and is shown as a cytoskeleton-associated mechanism that regulates tight junction assembly and function^87^. We also identified some areas where the TJ proteins seem to be missing between cells (highlighted with white arrows in Fig. 4).

In order to ascertain whether the redistribution and dislocation of tight junctional proteins is a synergistic effect of *C. jejuni* and TMAO on the Caco-2 cells, or solely form TMAO treatment, we conducted a similar experiment but in an uninfected model of Caco-2 cells. Specifically, we probed for occludin (Fig. 5A), ZO-1 (Fig. 5B), and ZO-3 (Fig. 5C). The IF images elucidated the sites of tight junctional complex with ruffling, and punctation highlighted with white arrows (Fig. 5). In addition, TMAO diminished the abundance of occludin and ZO-1 proteins at the membrane junctions (Fig. 5A,B). In addition, TMAO stimulated the redistribution of ZO-3 from the membrane to the cytoplasm (Fig. 5C). Taken together, these results suggest that the TMAO treatment of Caco-2 cells disrupts the functional localization of tight junction proteins.

### TMAO modulates the abundance of tight junctional proteins in the intestinal epithelial cells

The modulation of TJ proteins and the regulation of TJ complex by microbial products have been studied in the context of strengthening and weakening the intestinal barrier function and permeability^77,84,88^. Upon uncovering that TMAO promotes *C. jejuni* invasion in Caco-2 cells (Fig. 3), as well as it stimulates *C. jejuni* to disrupt the function of TJ proteins *in vitro* (Fig. 4 and Fig. 5), we further examined their level of total protein expression of the TJ proteins in the intestinal epithelial monolayer without *C. jejuni* infection (Fig. 6A-F) or with *C. jejuni* infection (Fig. 6G-L). Unlike IF staining, which only shows the localization of specific proteins in cellular compartments, here we detected total protein levels using Western blot analysis. As explained in Fig. 5, we followed a similar timeline for this experiment. Briefly, we subjected confluent monolayers of Caco-2 cells with varied concentrations of TMAO for 48 hours, followed by a final 3-hour microaerobic incubation (without *C. jejuni* infection) before harvesting proteins for Western blot analysis (Fig. 6A-F). We probed for tight junctional proteins claudin-4, occludin, ZO-1, ZO-2, and ZO-3. Intriguingly, we found that TMAO treatment and the terminal microaerobic incubation of Caco-2 cells without *C. jejuni* infection (Fig. 6A-F) resulted in the upregulation of total protein levels of both occludin (Fig. 6C) and ZO-2 (Fig. 6E) at 500 µM TMAO treatment. Also, 1 mM TMAO treatment was sufficient in increasing the total occludin protein level in the Caco-2 monolayers, while ZO-2 was no longer increased. TMAO did not alter other TJ proteins tested in this condition. The heightened levels of total occludin protein content in these monolayers (Fig. 6C) did not provide increased protection of the cell (Fig. 3 and Fig. 4). Hence, this data highlights that although TMAO increased the level of total cellular occludin proteins (Fig. 6C), however, it also displaced or redistributed occludin from the membrane TJ complex (Fig. 5A) possibly to cytoplasm. Together, these observations show that TMAO modulates the abundance of occludin and ZO-2 TJ protein levels in the intestinal epithelial cells without *C. jejuni* infection.

**Figure 6.**
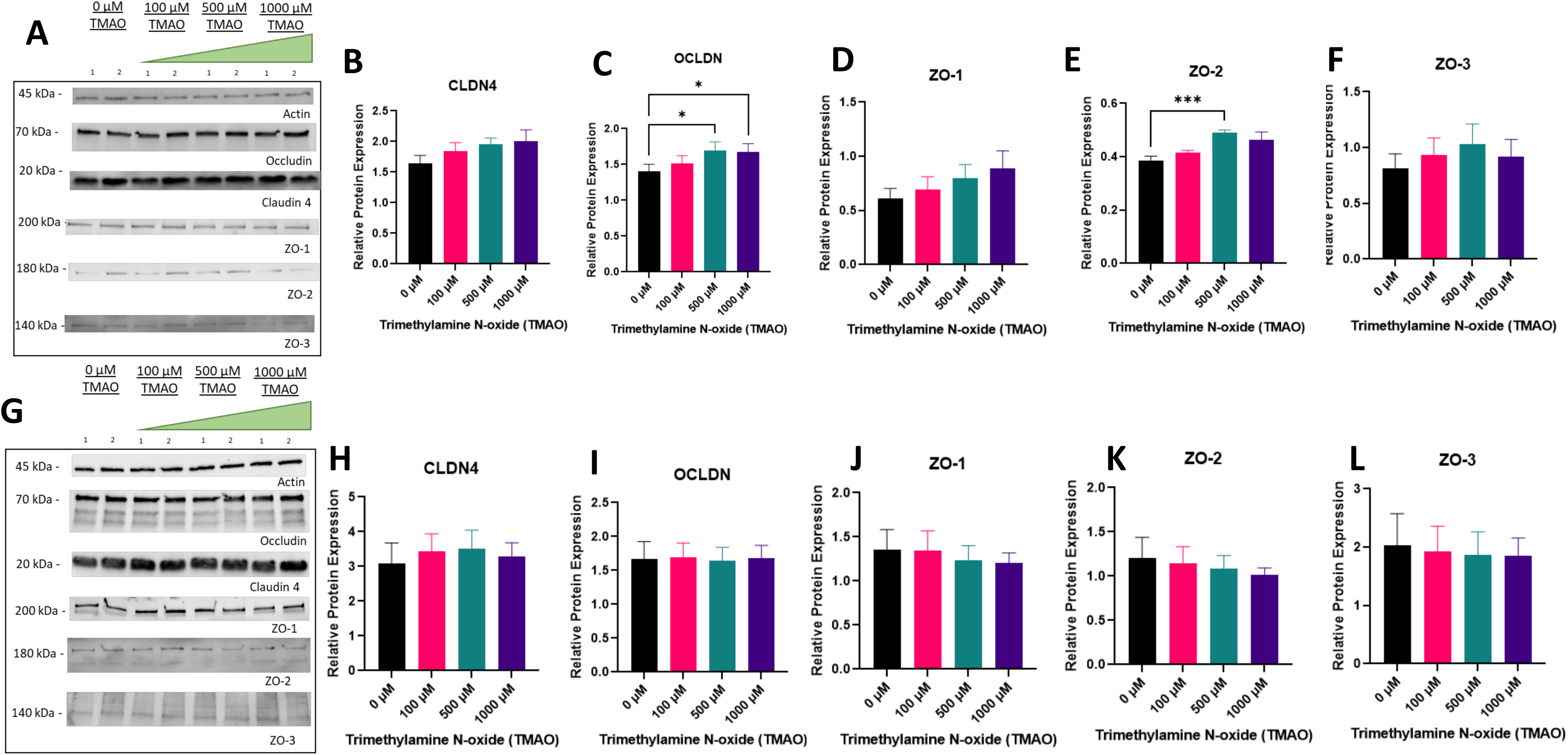
TMAO alters expression of total protein levels of occludin and ZO-2 without infection. (A) Representative images of immunoblotting of Caco-2 cells treated with TMAO for 48 hours at varied concentrations, with a terminal microaerobic incubation of 3 hours prior to protein harvesting. Quantification of (B) claudin 4 (CLDN4), (C) occludin (OCLDN), (D) zonula occludens 1 (ZO-1), (E) zonula occludens 2 (ZO-2), and (F) zonula occludens 3 (ZO-3). (G) Representative images of immunoblotting of Caco-2 cells treated with TMAO for 48 hours at varied concentrations, with a terminal infection with *C. jejuni* under microaerobic conditions, lasting 3 hours prior to protein harvesting. Quantification of (H) CLDN4, (I) OCLDN, (J) ZO-1, (K) ZO-2 and (L) ZO-3. Data are presented as the mean ± SEM of three independent biological replicates. Statistical significance was assessed via One-Way RM ANOVA followed by Dunnett’s post hoc test. * p<0.05, *** p<0.001

Next, we also evaluated the expression of these tight junctional proteins during TMAO treatment followed by *C. jejuni* infection in a microaerobic condition (Fig. 6G-L). As previously described for invasion assays (Fig. 3 & Fig. 4), we treated confluent monolayers of Caco-2 cells with varied concentrations of TMAO for 48 hours coupled with a terminal 3-hour microaerobic incubation with *C. jejuni* infection before proteins were harvested for western blot analysis. As demonstrated in Fig. 6G, we found the presence of multiple, smaller sized bands that were not present in the uninfected cells (compared to Fig. 6A). This banding pattern is expected as *C. jejuni* utilizes a secreted serine protease, HtrA, that cleaves occludin during infection and invasion. Further studies with HtrA-deficient *C. jejuni* strains are needed to confirm this observation. In addition, Fig. 6G-L demonstrated that *C. jejuni* infection abrogated the increases of total occludin protein (∼65 kDa band) (Fig. 6I vs. Fig. 6C) and ZO-2 (Fig. 6K vs. Fig. 6E) that was previously demonstrated by TMAO only treatment (Fig. 6C,E). *C. jejuni* seems to be capable of causing enough occludin cleavage during infection to reduce the total occludin protein increase stimulated by TMAO in these Caco-2 cells.

We then assessed the influence of TMAO on the Caco-2 cells under standard incubation conditions (95% Air, 5% CO2). In these assays, we treated confluent monolayers of Caco-2 cells with varied concentrations of TMAO for 48 hours. However, we omitted the final 3-hour microaerobic incubation and did not conduct infection with *C. jejuni*. We detected total protein levels using Western blot analysis. As shown in Fig. S3A-F, TMAO treatment showed no significant increase in any tested tight junctional complex proteins (claudin-4, occludin, ZO-1, ZO-2, ZO-3).

We further sought to investigate whether TMAO treatment and infection with *C. jejuni* cause transcriptional alterations of the genes encoding tight junction proteins. In these experiments, we maintained the identical timeline of cellular growth of Caco-2 cell monolayers (as in Fig. 3 & Fig. 4), including our two conditions: 1) TMAO-treated cells with a final 3-hour microaerobic incubation (Fig. 7A-E), and 2) TMAO-treated cells with a final *C. jejuni* infection under microaerobic conditions for 3 hours (Fig. 7F-J). We determined that there was no significant change in the transcriptional levels of any tested tight junctional complex proteins (claudin-2, claudin-4, ZO-1, ZO-2, ZO-3). These data show that neither TMAO treatment alone nor a 3-hour *C. jejuni* infection alters the transcriptional expression of tight junctional complex proteins.

**Figure 7.**
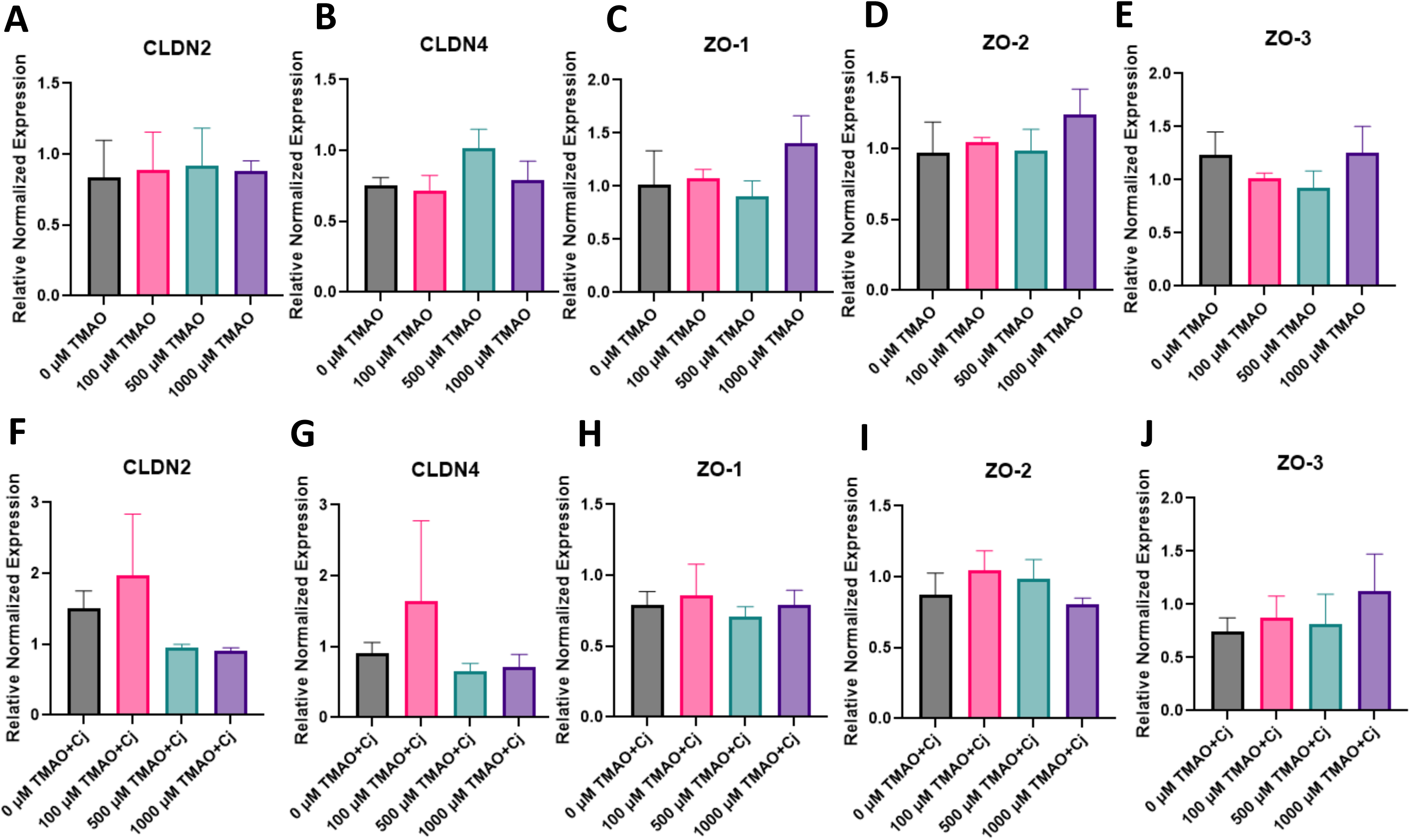
Relative mRNA expression levels of tight junction genes are not affected by TMAO treatment and *C. jejuni* infection. RT-qPCR of Caco-2 cells treated with TMAO for 48 hours at varied concentrations, with a final microaerobic incubation (with vehicle only, no *C. jejuni*) of 3 hours prior to RNA harvesting. (A) claudin 2 (CLDN2), (B) claudin 4 (CLDN4), (C) zonula occludens 1 (ZO-1), (D) zonula occludens 2 (ZO-2), and (E) zonula occludens 3 (ZO-3). RT-qPCR of Caco-2 cells treated with TMAO for 48 hours at varied concentrations, with 3 hours of *C. jejuni* infection under microaerobic conditions. (F) claudin-2 (G) claudin-4, (H) ZO-1, (I) ZO-2, and (J) ZO-3. Data are presented as the mean ± SEM of three independent biological replicates. Statistical significance was assessed via One-Way ANOVA followed by Dunnett’s post hoc test.

### TMAO synergistically reduces epithelial TEER and escalates *C. jejuni* paracellular transmigration

*C. jejuni* can invade intestinal epithelial cells during infection^1,74,78^, but also transmigrate across the intestinal epithelium, invade basolaterally and disseminate to other organs^79–82^. To investigate the effect of TMAO on *C. jejuni* paracellular transmigration, we utilized polarized Caco-2 cells. Cells were grown for both 14 and 21 days to allow the growth of Caco-2 cells to become a confluent and polarized monolayer on Transwell inserts. The transepithelial electrical resistance (TEER) of the monolayers increased over the 14 days (Fig. S4A) or 21 days (Fig. 8A) to 300 – 400 Ohm × cm2, representing the increases in the barrier maturation and proper polarization. The polarized monolayers were treated with 1 mM TMAO 48 hours before the final *C. jejuni* infection under microaerobic conditions for 3 hours. We measured the pre- and post-infection TEER values of the monolayers and bacterial transmigration of *C. jejuni* (Fig. 8A-C).

**Figure 8.**
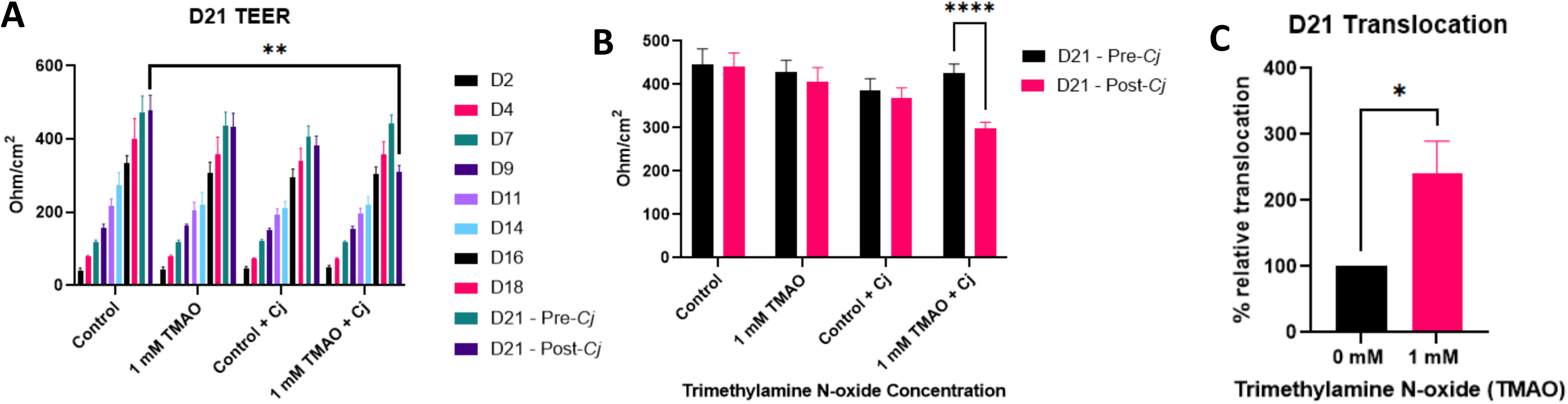
TMAO promotes transmigration of *C. jejuni* in the polarized Caco-2 cells. (A) The transepithelial electrical resistance (TEER) was measured across the polarized epithelial layer over 21 days. (B) The graph shows the TEER on day 21 before and after infection with *C. jejuni* in a microaerobic condition. (C) The relative transmigration of *C. jejuni* 81-176 across the polarized monolayer after *C. jejuni* infection for 3 hours in a microaerobic condition. Data are presented as the mean ± SEM of three independent biological replicates. Statistical significance was assessed via (A) Two-Way RM ANOVA followed by Dunnett’s post hoc test, (B) Two-Way RM ANOVA followed by Sidak’s post hoc test, and (C) Student’s t-test. * p<0.05 ** p<0.01, **** p<0.0001

First, we investigated the transmigration of *C. jejuni* in 14-day-old polarized epithelial cells. As shown in Fig. S4A-B, we did not find any significant differences in TEER values between the pre- and post-vehicle treatment (without *C. jejuni* infection) in 1 mM TMAO-treated wells. However, *C. jejuni* infection significantly reduced TEER. Interestingly, *C. jejuni* infection in the presence of TMAO remarkably lowered the TEER, indicating a dysfunction of epithelial integrity caused by *C. jejuni* and TMAO (Fig. S4B). Nonetheless, upon evaluation of bacterial transmigration (Fig. S4C), we found that TMAO treatment did not significantly increase the number of recovered *C. jejuni* (CFU) from the bottom chambers compared to the untreated wells.

We postulated that this timeline (14 days) was not sufficient to allow the full maturation of the barrier function of the polarized monolayer of the Caco-2 cell. Hence, we further examined the 21-day-old polarized monolayers (Fig. 8B). Interestingly, we did not find any significant perturbations in TEER values compared between pre- and post-microaerobic incubation of both the untreated and 1 mM TMAO-treated 21-day-old polarized monolayers. In addition, when polarized cells were infected with *C. jejuni* alone in the absence of TMAO, we found that *C. jejuni* alone did not reduce the TEER values between pre- and post-infection (Fig. 8B). However, when *C. jejuni* infected the polarized Caco-2 monolayer in the presence of TMAO, a significant drop in the TEER value was observed after the infection (Fig. 8B). These results indicate that *C. jejuni* causes significantly more epithelial barrier damage to the polarized Caco-2 monolayer in the presence of TMAO.

We next investigated the paracellular transmigration of *C. jejuni* in the 21-day-old polarized Caco-2 cells on the Transwell system (Fig. 8C). Following the infection, we cultured the serially diluted media from the bottom chambers and enumerated CFU. Intriguingly, our results revealed a significantly higher recovery of *C. jejuni* when infected in the presence of TMAO compared to the untreated control (without TMAO). Collectively, these data demonstrate that TMAO promotes *C. jejuni*-mediated dysfunction of the epithelial barrier and enhances the transmigration of *C. jejuni* across polarized colonocytes.

### TMAO enhances *C. jejuni*-mediated inflammatory response in Caco-2 cell monolayers

During infection, *C. jejuni* triggers the production of inflammatory cytokines, including IL-1β, IL-8, and TNFα in colonic epithelial cells^27,89^. To investigate the potential of TMAO to exacerbate the inflammatory response during *C. jejuni* infection in Caco-2 cells, we initially performed RT-qPCR assays targeting IL-1β, IL-8, and TNFα after 3 hours of infection with *C. jejuni*. Furthermore, we also examined infection durations for 4 hours and 6 hours. Our qPCR analysis did not reveal any significant differences in the gene expression of these three cytokines in response to TMAO treatment and *C. jejuni* infection for 3- and 4-hours (Fig. S5AB). However, we found significant alterations in IL-1β and IL-8 after 6 hours of infection in comparison to the untreated, uninfected control (Fig. S5G,H)

Finally, we performed RT-qPCR assays targeting IL-1β, IL-8, and TNFα after a 24-hour infection with *C. jejuni*. We found that TMAO alone did not stimulate the expression of cytokine genes, including IL-1β, IL-8, and TNFα. However, TMAO enhanced *C. jejuni*-mediated inflammatory response in the Caco-2 cells by stimulating expression of IL-1β, IL-8, and TNFα in comparison to the untreated, uninfected control (Fig. 9A-C). Furthermore, when we compared the expression of IL-1β and IL-8 genes in Caco-2 cells solely infected with *C. jejuni* alone in the absence of TMAO, we found the 1 mM TMAO synergistically heightened expression of these cytokine genes in the Caco-2 cells infected with *C. jejuni* and treated with TMAO (Fig. 9A,B). In addition, *C. jejuni* alone induced a significant expression of TNFα (Fig. 9C); however, 500 μM TMAO and *C. jejuni* infection surprisingly diminished the TNFα gene expression. These findings collectively suggest a synergistic effect of TMAO on *C. jejuni*-mediated upregulation of the expression of IL-1β and IL-8 cytokine genes.

**Figure 9.**
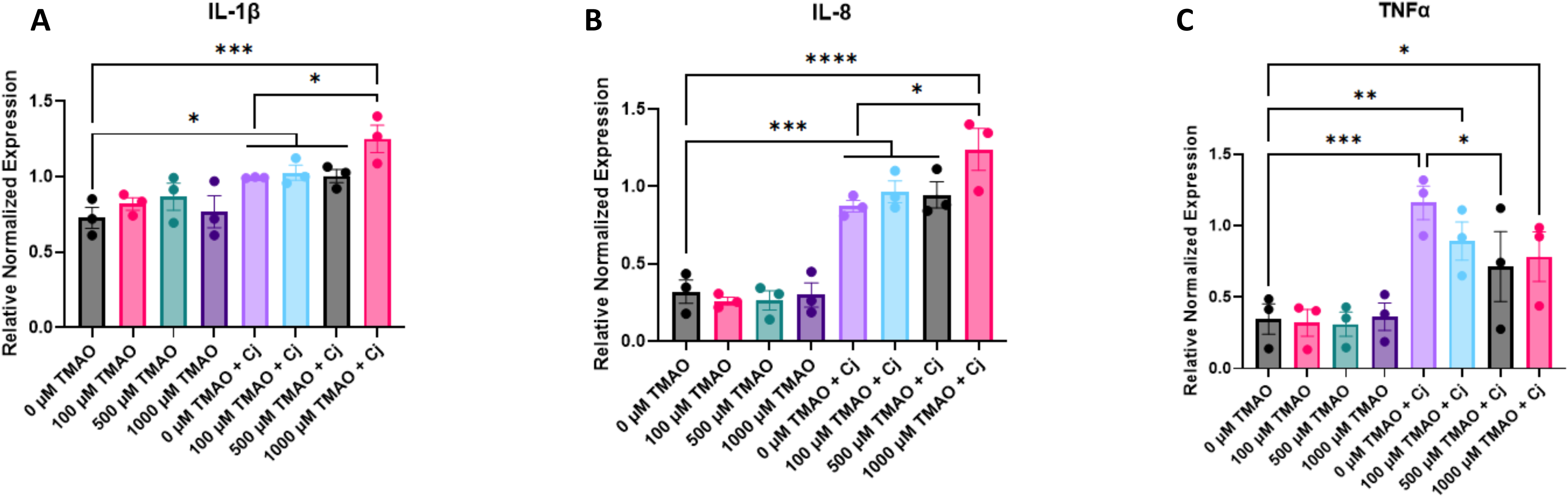
*C. jejuni* and TMAO synergistically increase the inflammatory response in Caco-2 cell monolayers. RT-qPCR of (A) IL-1β, (B) IL-8 and (C) TNFα of Caco-2 cells treated with TMAO for 48 hours at varied concentrations, and infected with *C. jejuni* for 24 hours. Monolayers were treated with TMAO and had either a final 4-hour incubation with vehicle under microaerobic conditions or were infected with *C. jejuni* for 4 hours under microaerobic conditions. After those 4 hours, media in all wells were replaced with fresh media supplemented with 20 ug/mL gentamicin for 18 hours, under 5% CO2. Data are presented as the mean ± SEM of three independent biological replicates. Statistical significance was assessed via One-Way ANOVA followed by Sidak’s post hoc test. * p<0.05, ** p<0.01, *** p<0.001, **** p<0.0001

Furthermore, we also examined shorter infection durations, including 3 hours, 4 hours, and 6 hours. Our qPCR analysis did not reveal any significant differences in the gene expression of these three cytokines in response to TMAO treatment and *C. jejuni* infection for 3- and 4-hours (Fig. S5AB). However, we found significant alterations in IL-1β and IL-8 after 6 hours of infection (Fig. S5C).

## DISCUSSION

As the complete mechanistic understanding of *C. jejuni*’s role in gastroenteritis remains elusive, this pathogen continues to pose a major health threat to humans, for its elevated rates of outbreaks^89–91^, post-infection sequalae^92^, and increasing antibiotic resistances^93^. A major limitation into studying *C. jejuni*’s pathogenesis and virulence factors remains a lack of infectible murine model, with the most commonly used model being an immuno-defective transgenic mouse model, the IL-10^-/-^ knockout strain^23^. The C57Bl/6 mouse demonstrates colonization resistance to *C. jejuni*, as the native gut microbiota in these mice raised under specific pathogen free conditions effectively inhibits pathogenic colonization. Investigating the attributes of colonization resistance in C57Bl/6 mice has been of increasing interest in the recent literature^24^. The gut microbiota confers colonization resistances through various methods, including modulation of the immune system, competitive exclusion, the production of antimicrobial molecules, and the production of microbial metabolites^94^. Microbial metabolites such as short chain fatty acids (SCFAs) and secondary bile acids have been studied more extensively than other microbial metabolites for their protective effects. In many reports, there has been a clear association with dysbiosis and the susceptibility and disease severity of multiple different enteric pathogens. For example, a ‘Western Diet’ (high protein and high fat) caused dysbiosis within a mouse model for increased the relative abundance of the pro-inflammatory Proteobacteria phylum, a decrease in protective SCFAs produced by the intestinal microbiota, and these conditions increased the susceptibility of mice to Adherent-Invasive *E. coli* infections^95^. Furthermore, the Heimesaat lab has also elucidated an association between a ‘Western-style’ diet and *C. jejuni* infection^55^, to which found increased Proteobacteria and *E. coli* levels in the gut microbiota of these mice. Further interested in elucidating this association between *C. jejuni* and intestinal *E. coli* levels, the Heimesaat lab found that exogenous supplementation of *E. coli* to specific-pathogen-free raised C57Bl/6 mice was capable of rendering these previously resistant mice susceptible to *C. jejuni* infection^56^. This association between *C. jejuni* infection and *E. coli* was later corroborated with a human study, showing that Poultry workers with increased intestinal levels of *Escherichia* had increased incidence of *C. jejuni* infection in comparison to their coworkers^54^, though no mechanism of how *E. coli* increases *C. jejuni* infection is not known.

In our study, we employed the well-established IL10KO mouse model for *C. jejuni* infection (Fig. 1). Following 2 weeks of infection with *C. jejuni*, we observed a marked increase in Proteobacteria and *E. coli.* Furthermore, the IL10KO mice were much more susceptible to *C. jejuni* infections compared to the WT mice, consistent with reported findings in the literature^23^. Our microbial community analysis demonstrated significant alterations in the composition of intestinal microbiota. Based on these findings, we further hypothesized that the microbial metabolites would have a critical role in *C. jejuni* infection process. Interestingly, we determined an increased abundance of TMAO in the *C. jejuni*-infected IL10KO mouse guts. Intriguingly, we also determined that *C. jejuni* infection dramatically increased *E. coli*’s choline TMA lyase *cutC* gene in the IL10KO mice colon but not in other groups of mice. Recently, it has been reported that some intestinal microorganisms, including *E. coli,* anaerobically metabolize diet-derived choline to TMA by choline TMA-lyase (cutC). Furthermore, TMA is oxidized to TMAO by flavin-dependent monooxygenase 3 (FMO3) in the host liver^38,40^. BLAST search and *C. jejuni* genome analysis confirmed that *C. jejuni* 81-176 does not encode a homologous gene to *E. coli*’s *cutC* gene. However, the *C. jejuni* mRNA encoding the TorA enzyme for TMAO reduction is also significantly elevated in *C. jejuni*-infected IL10KO mice. This observation underscores the significance of elevated levels of *E. coli* and TMAO metabolites during *C. jejuni* infection of IL10KO mice. Based on these findings, we thus postulate that the expansion of *E. coli* and its cutC gene in the infected IL10KO mice resulted in the increased abundance of TMAO, which promotes *C. jejuni* invasion, paracellular transmigration, barrier damage, and inflammatory response.

There is a growing recognition of the pivotal role the gut microbial metabolites play in enteric infections^24,96^. For example, in response to acid stress, many Proteobacteria species can produce polyamines to modulate the pH of the luminal environment, which can out-compete pathogens^96^. In addition, ingested dietary fibers can be metabolized by gut bacteria to produce protective SCFAs in the intestine. The SCFAs contribute to the maintenance of the anaerobic conditions within the intestinal lumen and also can impede infection by affecting metabolic function and the intracellular pH of pathogenic bacteria^24^. Previously it was reported that bile acid induces expression of type three secretion system genes in a non-O1/non-O139 *Vibrio cholerae* strain and promotes its pathogenesis by regulating ToxR/VttR regulatory network^97^. In contrast, many primary and secondary bile acids have antimicrobial properties, and therefore, the secondary bile acids producing bacteria in the intestinal tract are associated with providing colonization resistance^24^.

TMAO is a microbial metabolite that is implicated in multiple different chronic inflammatory diseases, including but not limited to kidney diseases, metabolic syndrome and related conditions, cardiovascular disease, and cancer^40^. TMAO causes platelet hyperreactivity, foam cell formation and adhesion, cardiac fibroblast migration and proliferation, and cellular apoptosis that all contribute to cardiovascular disease progression. TMAO exacerbates the inflammatory response by increasing MCP-1, TNF, IL-6, IL-1, IL-18 and the activation of the NLRP3 inflammasome in kidney disease^98^. In Metabolic Syndrome related diseases, TMAO was implicated in causing endoplasmic reticulum stress and insulin resistance in patients at risk for diabetes^99^, and also implicated in promoting de novo lipogenesis in the liver of metabolic-dysfunction-associated fatty liver disease^100^. Increased serum levels of TMAO in colorectal cancer are correlated to increased tumors, worse stage and incidence of metastasis and overall lower disease survival^40^. Increasing amount of studies are also highlighting that TMAO may be linked to other cancers as well, such breast, ovarian and gastric cancers^101^. Yet, the role of TMAO has not been fully mechanistically determined in the progression or severity of many diseases, with its association primarily established through serum levels and disease prognosis rather than through detailed mechanistic understanding.

*C. jejuni* utilizes TMAO as a terminal electron acceptor in environments with limited oxygen availability, such as the conditions found within the anaerobic colonic lumen during infection, via the functions of TMAO Reductase TorA^49^. Nevertheless, the precise involvement of TMAO, especially its role in epithelial cell invasion, paracellular transmigration and chemotaxis remained unclear. Here we present novel findings for the first time that *C. jejuni* can perceive TMAO, along with its precursor molecules, including L-carnitine and betaine, acting as chemoeffectors. Furthermore, our studies confirmed TMAO’s role as a chemoattractant for *C. jejuni*. Having upwards of 25 different methyl accepting chemotaxis proteins, *Campylobacter* species includes systems for energy taxis and aero taxis, nearly all of which have multiple different chemoeffectors that can be sensed and induce a response in the bacterium. As an alternative electron acceptor, it would be intriguing to investigate further how *C. jejuni* can perceive TMAO with its energy taxis chemoreceptor system^5^. Chemotaxis is a necessary virulence factor of *C. jejuni*, as genetic mutations and deletions of different *C. jejuni* methyl-accepting chemotaxis proteins, have resulted in decreased adherence and invasion *in vitro*^31,63^, and decreased colonization and infection in chicken ^31,32^ and mouse models^30^. When comparing to other microbial metabolites, SCFAs within the colonic lumen, such as butyrate and acetate increased *C. jejuni* virulence and supported colonization within the host colon by modulating *C. jejuni* amino acid catabolism and amino acid transport^6,102^. Interestingly, L-lactate represses *C. jejuni* colonization within the upper intestinal tract^6^, however, paradoxically, L-lactate promotes *C. jejuni* growth and expansion in the inflamed colon, as well as increased invasion and adherence *in vitro*^103^. The short chain fatty acid butyrate is a metabolite commonly produced by commensal organisms, which negatively impacted *C. jejuni* motility^104^. *C. jejuni* also responds to bile acids within the intestinal tract, which causes increased outer membrane vesicle production, and overall increased virulence of *C. jejuni*^25,26^. In addition, bile acids are usually chemorepellents and are sensed by Tlp3 and Tlp4. Hugdahl et al provided evidence that bovine and ox biles function as chemoattractants; intriguingly purified bile acids, including cholic acid and deoxycholic acids were found to be chemorepellent^105–107^.

The use of TMAO as an alternative electron acceptor in *C. jejuni* has been characterized to cause transcription regulation changes via the RacRS two-component system^68^. From this information and the increase of TMAO in our IL10KO mouse model, we further postulate that the utilization of alternative respiration pathways would increase *C. jejuni* virulence by enabling survival and providing energy production in a diverse array of microenvironments. Understanding the connection between *C. jejuni* and TMAO, we investigated further the mechanistic role TMAO plays in pathogenic infection. First, we determined that the presence of TMAO was sufficient in causing an increased invasion of *C. jejuni* into Caco-2 cell monolayers, and this effect was not mediated via toxin production (Fig. 2). We also further elucidated that *C. jejuni* caused increased intestinal barrier damage of polarized Caco-2 cell monolayers, directly correlating to increased transmigration of pathogen across that monolayer (Fig. 8). When comparing to other pathogens in response to TMAO, *V. cholerae* had increased *in vivo* colonization and increased mortality rate of mice when infected in a suspension containing TMAO^46,108^, and uropathogenic *E. coli* also had increased colonization of bladder epithelial cells in the presence of TMAO *in vitro*^48^. Our results reciprocate the capability of TMAO to increase the virulence and colonization of pathogenic bacteria.

The effects of TMAO on intestinal epithelial cells and their intestinal barrier and tight junctional function have not yet before been elucidated, while *C. jejuni* has been characterized to cleave tight and adherens junctions’ proteins to cross the intestinal epithelial layer^109^. Here, we determined that TMAO causes increased levels of tight junctional proteins occludin and zonula occludens 2 (Fig. 6). These protein increases were abrogated upon infection with *C. jejuni,* potentially due to the cleavage of occludin during infection by this pathogen. Though we determined no effect on the transcriptional levels of proteins claudin-2, claudin-4, ZO-1, ZO-2, or ZO-3, we saw in immunofluorescence staining that TMAO treatment induced disruptions to tight junctional complexes, for more jagged, ruffled, and punctuated cell-to-cell junctions. We determined this effect for occludin, ZO-1, and ZO-3 (Fig. 4, Fig. 5). Remarkably, we found that *C. jejuni* diminished the expression of claudin-4 independent of TMAO (Fig. S2C). From these results, we determined that TMAO-treatment alone caused the dissociation of tight junctional complex proteins from the cell-to-cell junctions, disrupting the overall function of the tight junctional complex proteins.

As a microbial metabolite, TMAO has also been implicated as a biomarker in many inflammatory mediated diseases, as mentioned previously. Considering the inflammatory response, we found that after the 48-hour treatment of Caco-2 cells with TMAO had no significant effect on IL-1β, IL-8 or TNFα expression (Fig. 9, Fig. S5). Alternatively, we determined that the 48-hour treatment of Caco-2 cells and presence of TMAO during *C. jejuni* infection synergistically increased IL-1β and IL-8 after 6 and 24 hours of infection. However, C. jejuni did not induce cytokine expression when infection was conducted for a shorter time of 3 and 4 hours. The synergistic effect of TMAO and other pathogens has been reported previously. In the case of *Vibrio cholerae,* TMAO incubation increased cholera toxin production under anaerobic conditions, such as those found within the colonic lumen, in a stringent response to the growth inhibition that occurs in anaerobic conditions^46,108^, and this was also correlated to the ability of *V. cholerae* to colonize *in vivo*. *Helicobacter pylori* synergistically increased the inflammatory response of gastric epithelial cells *in vitro* in the presence of TMAO, increasing IL-6 and CXCL1 protein and transcriptional levels^46,47^. For uropathogenic *Escherichia coli*, TMAO synergistically caused and increased colonization of bladder epithelial cells coupled with an increased inflammatory response, specifically increasing the levels of IL-1β, IL-6, Il-8, CXCL1, and CXCL6^47^. Our results recapitulate these data, as *C. jejuni* has increased invasion, transmigration, epithelial damage of Caco-2 cells *in vitro*, as well as increasing the inflammatory response of these monolayers, via increased IL-1β and IL-8 transcripts. Inflammatory cytokine is a critical regulator of TJ function. Our data demonstrated that during early time points of *C. jejuni* infection in Caco-2 cells (3 and 4 hours), there were no significant alterations in the expression of the cytokines. However, TMAO and *C. jejuni* synergistically enhanced cytokine production after 6 hours. These data suggest that TJ proteins’ dysfunction of the epithelial cells was potentially caused by bacterial factors and metabolites but probably not directly by cytokines’ action on the epithelial cells.

This research, as many others, is not without its caveats. Firstly, we would like to note that there are many components when researching the intestinal microbiota, which makes it difficult to isolate one bacterial species, such as commensal *E. coli*, as a determinant in *C. jejuni* infection. While this is true, there have also been multiple established labs that have already elucidated this correlation between commensal *E. coli* and *C. jejuni* susceptibility of infection, making it a prime target for future work. The same can be mentioned for investigating a single microbial metabolite. We are currently working towards elucidating other microbial metabolites that we determined with the global metabolomics analysis from our mouse model data, all of which are highly correlated to *E. coli* metabolism. While we have not fully teased out a mechanism for TMAO on *C. jejuni* pathogenesis, we are also currently working on elucidating the regulatory systems that TMAO may modulate in *C. jejuni*, as it has been previously determine that TMAO transcriptionally regulates other respiratory systems within *C. jejuni*.

To summarize, in this study, we uncovered that *C. jejuni* infection in IL10KO mice promotes elevated abundance of *E. coli*, its choline TMA lyase *cutC* gene, and its metabolite TMAO. We demonstrated that TMAO plays a pivotal role in exacerbating *C. jejuni* invasion, paracellular transmigration, and epithelial damage. Furthermore, we determined that TMAO promotes *C. jejuni* to increase the inflammatory immune response *in vitro*. These findings contribute to our understanding of the intricate interplay between *C. jejuni* and the intestinal microbiota, elucidate the influence of microbial metabolites on virulence, and aid in the overall understanding of *C. jejuni* pathogenesis.

## MATERIALS AND METHODS

### Bacterial Growth, Strains and Growth Curves

*Campylobacter jejuni* strain 81-176, as well as an isogenic insertional deletion mutant 81-176Δ*cdtABC*, were used in this study. Strains were grown on Brucella agar with (Hardy) or without blood (HI Media) for 48 hours at 42°C under microaerobic conditions, generated by GasPak EZ CampyPouch system (Becton, Dickson). For growth curve assays, *C. jejuni* was resuspended from plates into either control or 1.0 mM TMAO in MEME medium (Sigma), supplemented with 2 mM L-Glutamine (Gibco), 1% non-essential amino acids (Sigma), and 10% heat inactivated fetal bovine serum (Gibco) at an OD_600_ of 0.1 (∼10^7^ CFU). *C. jejuni* enumeration was conducted periodically over 21 hours via measurement of OD_600_ on the Nanodrop spectrophotometer.

### IL10KO Mouse Model

Both male and female 6-week-old 129Sv/Ev background WT and IL10KO mice were used in this study. Mice were maintained as per the approved protocol by the Institutional Animal Care and Use Committee. Mice were maintained at an average temperature of 22° C and a standard 12-hour light/dark cycle (7 am – 7 pm) and mice were fed regular chow *ad libitum*. After a week acclimation, the mice were gavaged with 0.3 mL 5×10^9^ CFU of *C. jejuni* strain 81-176 in PBS, or 1x PBS as a control for two consecutive days. These mice were monitored for two weeks for symptoms of infection, while stool was collected for 16s rRNA sequencing and CFU determination of *C. jejuni* during infection. Afterwards, the mice were euthanized, and the colon and luminal contents were collected. The luminal contents of the colon were collected for metabolomic analysis before the colonic tissues were processed for further analysis.

### 16S rRNA Sequencing

DNA was extracted from luminal contents using the Power Soil kit from MoBio Laboratories. To amplify the 16S rRNA genes, a composite forward primer and a reverse primer containing a unique 12-base barcode were used^7,53,110^. These barcodes were designed using the Golay error-correcting scheme to tag PCR products from respective samples. For paired-end 16S community sequencing on the Illumina platform, we used primers specific for the V4 region of the 16S rRNA gene, namely bacteria/archaeal primer 515F/806R. The forward PCR primer sequence contained the sequence for the 5′ Illumina adapter, the forward primer pad, the forward primer linker, and the forward primer sequence. Each reverse PCR primer sequence contained the reverse complement of the 3′ Illumina adapter, the Golay barcode (each sequence containing a different barcode), the reverse primer pad, the reverse primer linker (CC), and the reverse primer. The sequences of these primers have been published. For each sample, three independent PCR reactions were performed, combined, and purified with Ampure magnetic purification beads (Agencourt). The resulting products were visualized and quantified using gel electrophoresis, and a master DNA pool was generated from the purified products in equimolar ratios. The pooled products were then sequenced using an Illumina Miseq sequencing platform. Sequences were assigned to operational taxonomic units (OTUs) with UPARSE using 97% pairwise identity and were classified taxonomically using the RDP classifier retrained with Greengenes. After removing chimeras, the average number of reads per sample was 28,432. A single representative sequence for each OTU was aligned using PyNAST, and a phylogenetic tree was built using FastTree. The phylogenetic tree was used to compute the UniFrac distances.

### Determination of Inflammatory Markers and Metabolome Profiling

Samples from mouse distal colon were harvested and stored at −80°C. Then thawed samples were extracted by agitation with 1mL of degassed acetonitrile: isopropanol: water (3:3:2) at –20°C after which the soluble portion were recovered by centrifugation. Aliquots (450 μL) of that supernatant were used for each LC-MS analysis. Specifically, we used an ultra-performance liquid chromatograph. We injected on reverse phase as well as normal phase columns and collected in both positive and negative modes. In addition, we collected ion fragmentation (MS2) spectral to aid in feature identification. Raw mass spectrometry spectral data were collected for each biological replicate at each time point both in positive and negative MS modes. The data were processed to report integrated areas of each detected peak in individual samples. For this study paired and unpaired non-parametric tests were carried out. Features were listed in a feature list table and as an interactive cloud plot, containing their integrated intensities (extracted ionchromatographic peak areas) observed fold changes across the sample groups. Identifications were made by comparing retention time and tandem MS fragmentation patterns to the sample and a standard compound. Stool and tissue samples were collected and homogenized for ELISA assays for inflammatory markers tumor necrosis factor α (TNF-α) and lipocalin-2 (LCN-2), using ELISA Kits (Abcam).

### Chemotaxis Assays

Plates for chemotaxis had a base of 20% Brucella Agar (HI Media #M1039), which gave a final concentration of 0.3% agar, standard of motility agar conditions. These plates were supplemented with chemoeffectors of interest in deionized water, or deionized H_2_O as a control, for a final concentration of 100 mM. In each plate, 50 mL of 20% brucella agar was poured into a 100×15 Petri dish (VWR) and allowed to solidify. For the assay, 10 uL of OD_600_ = 30 of *C. jejuni* was stabbed and spotted in 3 locations within the agar plate. These plates were placed at 37°C in microaerobic conditions using GasPak EZ CampyPouch system (Becton, Dickson) and imaged after 24 hours. The diameters of the rings of chemotaxis were measured for comparison. Chemoeffectors that were tested included a vector (no added chemoeffector) control, serine (Sigma), L-carnitine (Sigma), betaine (Sigma), and trimethylamine N-oxide (TMAO) (Sigma).

### Cell Culture Maintenance and Metabolite Treatments

Caco-2 cells (ATCC HTB-37) were routinely grown in T_75_ flasks (Corning), and media was replaced every 2-3 days with MEME medium (Sigma), supplemented with 2 mM L-Glutamine (Gibco), 1% non-essential amino acids (Sigma), penicillin-streptomycin (Sigma) and 10% heat inactivated fetal bovine serum (Gibco), and passaged upon 90% confluency. For treatment assays, Caco-2 cells were treated with trimethylamine N-oxide (Sigma) at varied concentrations for a total of 48 hours including subsequent assays. We also utilized antibiotic-free media, composed of MEME medium (Sigma), supplemented with 2 mM L-Glutamine (Gibco), 1% non-essential amino acids (Sigma), and 10% heat inactivated fetal bovine serum (Gibco), which was used to feed cells the day prior to TMAO treatment for all cells, regardless of *C. jejuni* infection or not. For treatment assays, Caco-2 cells were treated with trimethylamine N-oxide (Sigma) at varied concentrations for a total of 48 hours including subsequent assays.

### Invasion Assays

Caco-2 cells were seeded into 24-well plates (Thermo Fisher) at a density of 1.5×10^5^ cells/cm^2^ and grown for 4 days before treatment with TMAO at varied concentrations for the experiment. After 48 hours of treatment, these cells would be infected at a MOI of 200 of *C. jejuni* 81-176 WT or Δ*cdtABC*, resuspended in 1x Hanks buffered saline solution (HBSS) (Gibco) according to Liu et al.^16^. After 3 hours at 37°C, either in 5% CO_2_ or microaerobic (GasPak EZ CampyPouch system, Benton, Dickson), cells were washed with 1X HBSS, before the continuation of a gentamicin protection assay, as previously described^73,74^. MEME medium, supplemented with 2 mM L-Glutamine, 1% non-essential amino acids and 10% fetal bovine serum (Caco-2 media) supplemented with gentamicin (Sigma) for a final concentration of 250 ug/mL was added to each well. This gentamicin protection assay was 2 hours of incubation at 37°C in 5% CO_2_. After, cells were washed in HBSS, lysed with an experimentally determined 0.1% Triton-X 100 (Sigma) for 15 minutes, and serially diluted into HBSS for plating on Brucella Agar plates. After 48 hours of growth, the colonies forming units (CFU) were counted for determination of invasion.

### Immunofluorescence Staining

Caco-2 cells were grown and treated in the same conditions as invasion assays, seeded into Nunc Lab Tek 8-well Chamber Slides (Thermo Fisher) at a density of 1.5×10^5^ cells/cm^2^ and grown for 4 days before treatment with TMAO at varied concentrations for the experiment. After 48 hours of treatment, these cells would be infected at a MOI of 200 of *C. jejuni* 81-176 WT, resuspended in 1x HBSS. After 3 hours at 37°C, under microaerobic conditions, cells were washed with 1X HBSS, before the addition of Caco-2 cell culture media plus the addition of 250 ug/mL gentamicin (Sigma). This gentamicin protection assay was 2 hours of incubation at 37°C in 5% CO_2_. After, cells were washed in HBSS again, before the addition of 4% paraformaldehyde (Thermo Fisher) for 15 minutes for fixation. The cells were then permeabilized with 0.1% Triton-X 100 for 10 minutes. After permeabilization, the cells were blocked in a 3% BSA solution in tris-buffered saline with the addition of 0.1% tween-20 (TBST) for an hour at room temperature. The slide was then probed with an antibody according to the manufacturer’s instruction. The antibodies used in this manuscript: *C. jejuni* mAb (Novus Biologicals), *C. jejuni* pAb (Invitrogen), claudin-4 (Invitrogen), occludin (Cell Signaling Technologies), zonula occludens-1 (Cell Signaling Technologies), zonula occludens-2 (ZO-2) (Cell Signaling Technologies), zonula occludens-3 (ZO-3) (Cell Signaling Technologies), anti-mouse IgG (Alexa Fluor® 488) (Cell Signaling Technologies), anti-mouse IgG (Alexa Fluor® 555)( Cell Signaling Technologies), anti-rabbit IgG (Alexa Fluor® 488) (Cell Signaling Technologies), and anti-rabbit IgG (Alexa Fluor® 555) (Cell Signaling Technologies).

### Immunoblotting for Tight Junctional Proteins

Caco-2 cells were grown in the same fashion as for the invasion assays. Whole cell lysates were collected for cells solely treated with TMAO and cells treated with metabolite and infected with *C. jejuni* 81-176. For the growth, Caco-2 cells were seeded into 24-well plates (Thermo Fisher) at a density of 1.5×10^5^ cells/cm^2^ and grown for 4 days before treatment with TMAO at varied concentrations for the experiment. After 48 hours of treatment with and without a terminal 3-hour microaerobic incubation, uninfected cell lysates were collected. For infected cells, an invasion assay would be conducted with a MOI of 200 of *C. jejuni* 81-176 WT, resuspended from in 1x HBSS. After 3 hours at 37°C, either in 5% CO_2_ or microaerobic (GasPak EZ CampyPouch system, Becton, Dickson) cells were washed with 1X HBSS, before lysis. Cell lysis was conducted with RIPA Buffer (Sigma) supplemented with a phosphatase and protease inhibitor (Thermo Fisher). This cell lysis buffer was added to the wells, cells were scraped from the wells and placed into Eppendorf tubes. The cell lysate is then placed on a sample shaker for 20 minutes at 4°C before centrifugation at 12,000 RPM for 20 minutes at 4°C to collect the supernatant. Protein concentrations of samples were then determined with a MicroBCA (Thermo Fisher). These concentrations were used to load equal sample of proteins into Any-kD^TM^ Mini-PROTEAN® TGX^TM^ Precast Protein Gels (BioRad) for SDS-PAGE. After the gel electrophoresis, proteins were transferred to a nitrocellulose membrane (BioRad). Immunoblotting protocols were then followed according to company protocol for each antibody. The following antibodies were used: Actin (Invitrogen), claudin-4 (Invitrogen), occludin (Cell Signaling Technologies), zonula occludens-1 (Cell Signaling Technologies), zonula occludens-2 (ZO-2) (Cell Signaling Technologies), and zonula occludens-3 (ZO-3) (Cell Signaling Technologies).

### Reverse Transcriptase quantitative Polymerase Chain Reaction (RT-qPCR) for tight junctional complex proteins and inflammatory markers

Caco-2 cells were grown in the same fashion as for the invasion assays. Whole cell lysates were collected for cells solely treated with metabolites and cells treated with metabolite and infected with *C. jejuni* 81-176. For the growth, Caco-2 cells were seeded into 96-well plates (Corning) at a density of 1.5×10^5^ cells/cm^2^ and grown for 4 days before treatment with TMAO at varied concentrations for the experiment. After 48 hours of treatment with a terminal 3-hour microaerobic incubation, uninfected cell lysates were collected. For infected cells, an invasion assay would be conducted with an MOI of 200 of *C. jejuni* 81-176 WT, resuspended in 1x HBSS. For the different treatment conditions, the cells infected for 3 and 4 hours at 37°C under microaerobic conditions (GasPak EZ CampyPouch system, Becton, Dickson) were lysed immediately following incubation. All other longer incubations were infected for 4 hours at 37°C under microaerobic conditions (GasPak EZ CampyPouch system, Becton, Dickson) before the media was replaced with fresh Caco-2 cell media containing 20 μg/mL gentamicin for the remained of the incubation period. After the said infection incubations, cells were washed with 1X PBS, before lysis. Lysis was conducted utilizing the SingleShot Cell Lysis Kit (BioRad), with its defined protocol. These lysates were then used with the iTaq Universal SYBR Green One-Step Kit (BioRad) for RT-qPCR in the BioRad CFX96, C1000 Touch Thermo Cycler. We used the following PrimePCR primers from BioRad: interleukin-1β (IL-1β), interleukin-8 (Il-8), tumor necrosis factor α (TNF-α), claudin 2 (CLDN2), claudin 4 (CLDN4), zonula occludens-1 (TJP1), zonula occludens-2 (TJP2), zonula occludens-3 (TJP3) and reference gene RPS18. For Data Analysis, we utilized the CFX Maestro Program that accompanies our BioRad CFX96, C1000 Touch Thermo Cycler.

### Bacterial Transmigration and Intestinal Barrier Integrity Assays *in vitro*

Caco-2 cells were seeded into 24-well Transwell plates (VWR) at a density of 1.5×10^5^ cells/cm^2^ and grown at 37°C in 5% CO_2_ for 14 or 21 days for confluency. Media was replaced every 2-3 days. Immediately following media replacement, transepithelial electrical resistance (TEER) was measured in Ohms with an EVOM2 meter (World Precision Instruments), using STX2 chopstick electrodes (World Precision Instruments) of each monolayer, in comparison to an empty Transwell for a blank. On day 12 or day 19, cells were treated in only the top chamber with either vehicle (nf-H_2_O) or TMAO for a final concentration of 1 mM, for 48 hours. After treatment, cells were infected with *C. jejuni* 81-176 at a MOI of 200 and incubated for 3 hours at 37°C in microaerobic conditions (GasPak EZ CampyPouch system, Becton, Dickson). Pre- and Post-infection with *C. jejuni*, TEER measurements were taken. Then, the media from the bottom well was collected for serial dilution plating for CFU counting of transmigrated bacteria on Brucella agar plates, which were grown at 37°C for 48 hours under microaerobic conditions (GasPak EZ CampyPouch system, Becton, Dickson).

### Statistical Analysis

All data are expressed as mean ± SEM from three or more independent experiments, and the level of significance between two groups was assessed with Student’s t-test. For more than two groups, one-way ANOVA or two-way ANOVA was conducted, followed by a Fisher’s LSD, Dunnett, or Sidak post hoc test. A value of p<0.05 was considered to be statistically significant. All data was analyzed within GraphPad Prism 10 software with the most appropriate statistical test based on the experimental conditions. Details of statistics are within the figure legends.

## SUPPLEMENTAL MATERIAL

All supplemental data follows the references. For final submission, will have details listed as needed.

## ACKNOWLEDGEMENTS

We acknowledge Dr. Carol Pickett for the use of *C. jejuni* deletion mutant strain 81-176 Δ*cdtABC*. We’d like to also acknowledge the UKY Light Microscopy Core for use of their Confocal Microscope for imaging.

## SUPPLEMENTAL FIGURE LEGENDS

**Supplemental Figure 1.**
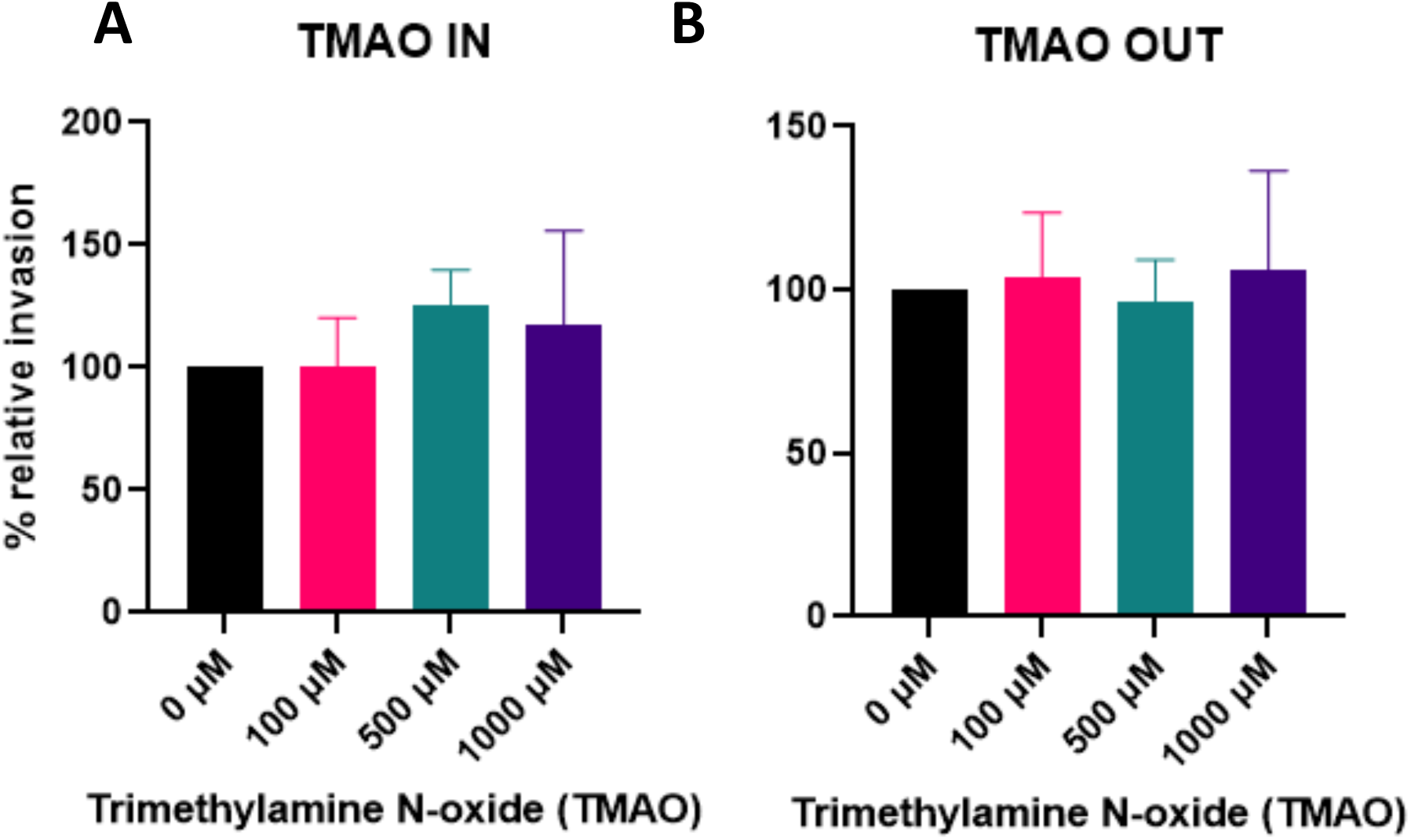
TMAO treatment nor presence had a result on *C. jejuni* invasion into Caco-2 cells under standard oxygen conditions. (A) Invasion Assay results of recoverable *C. jejuni* in Caco-2 cells treated for 48 hours with TMAO, with the metabolite left in the media during the infection step, under 5% CO_2_ conditions. (B) Invasion Assay results of recoverable *C. jejuni* in Caco-2 cells treated with TMAO for 48 hours before the removal of all media for conditioned media, eliminating the presence of TMAO during infection. Data are presented as the mean ± SEM of three independent biological replicates. Statistical significance was assessed via One-Way ANOVA followed by Fisher’s LSD post hoc test. ** p<0.01, **** p<0.0001

**Supplemental Figure 2.**
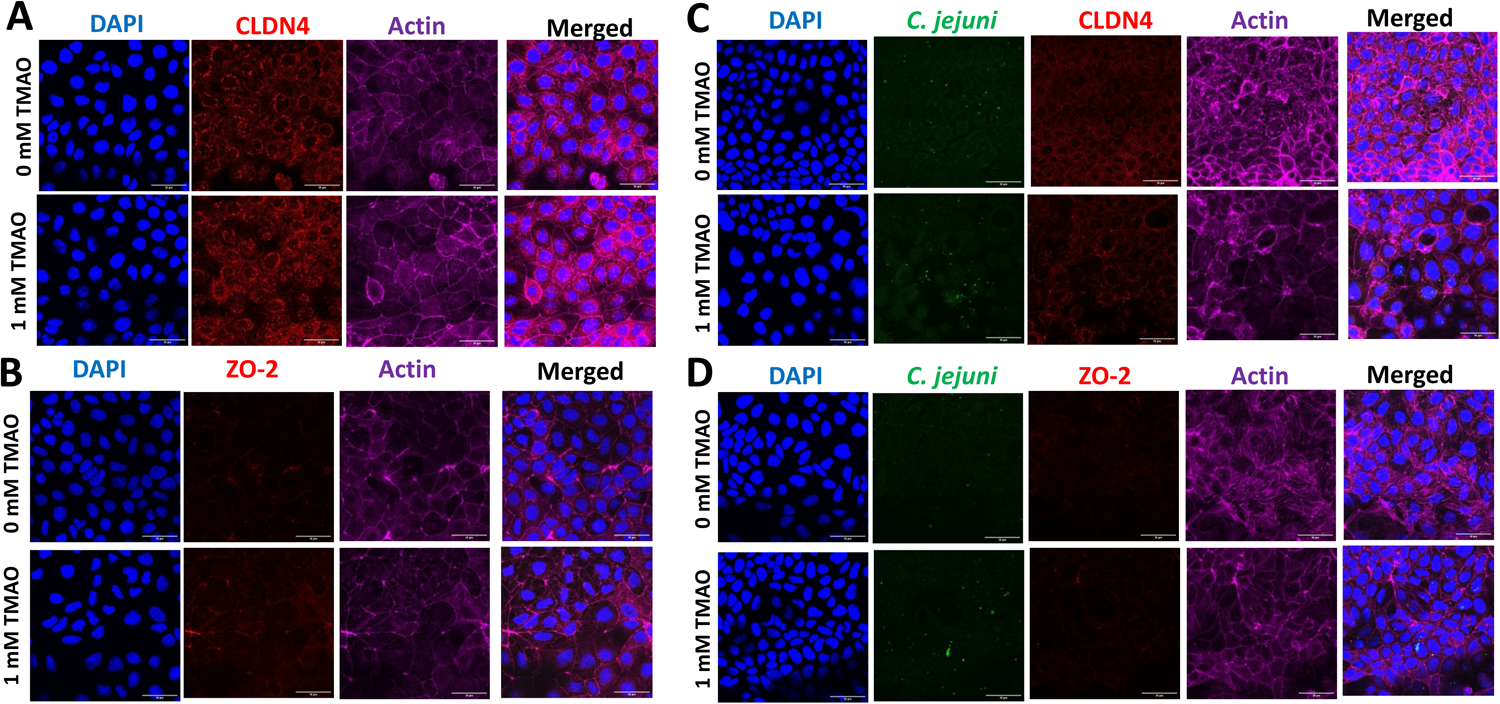
TMAO treatment, nor *C. jejuni* infection affect claudin-4 and ZO-2 function in Caco-2 cells. (A-D) Representative images from fixed cells and immunofluorescence staining of the nucleus (DAPI), tight junctional complex protein (red), Actin (purple), and *C. jejuni* (green). IF staining was conducted under 4 different experimental conditions: no-treatment control, TMAO-treated cells, *C. jejuni* infected control and *C. jejuni* infected TMAO-treated cells. Cells were treated with TMAO for 48 hours and the 3-hour infection with *C. jejuni* was conducted under microaerobic conditions, and uninfected conditions included a terminal 3-hour microaerobic incubation. Tight junctional complex proteins tested: (A, C) claudin 4 (CLDN4), and (B, D) zonula occludens-2 (ZO-2).

**Supplemental Figure 3.**
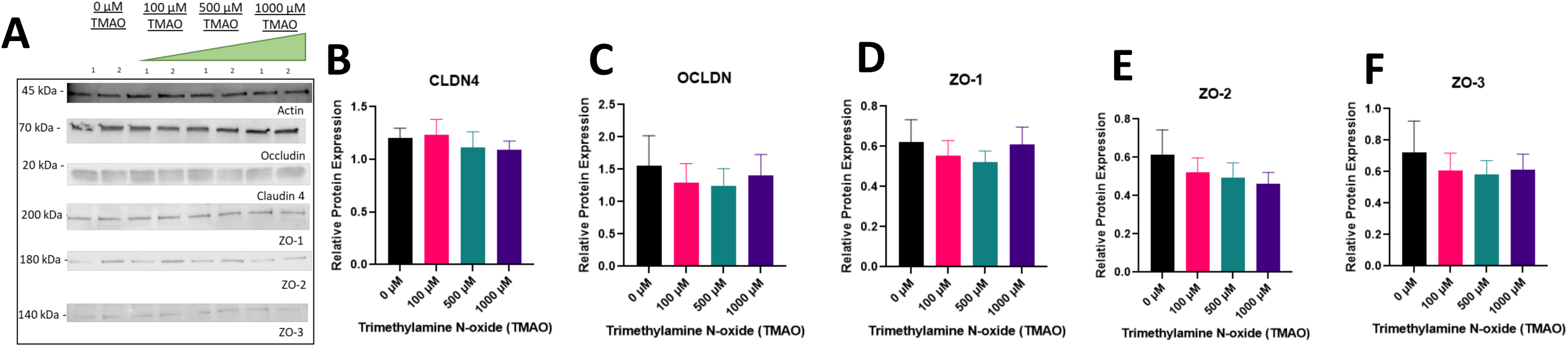
TMAO treatment of Caco-2 cells under standard oxygen conditions has no effect on protein levels of tight junctional complex proteins. (A) Representative images. Immunoblotting of Caco-2 cells treated with TMAO for 48 hours at varied concentrations prior to protein harvesting under standard oxygen conditions, of (B) claudin 4 (CLDN4), (C) occludin (OCLDN), (D) zonula occludens 1 (ZO-1), (E) zonula occludens 2 (ZO-2), and (F) zonula occludens 3 (ZO-3). Data are presented as the mean ± SEM of three independent biological replicates. Statistical significance was assessed via One-Way RM ANOVA followed by Dunnett’s post hoc test. * p<0.05, *** p<0.001

**Supplemental Figure 4.**
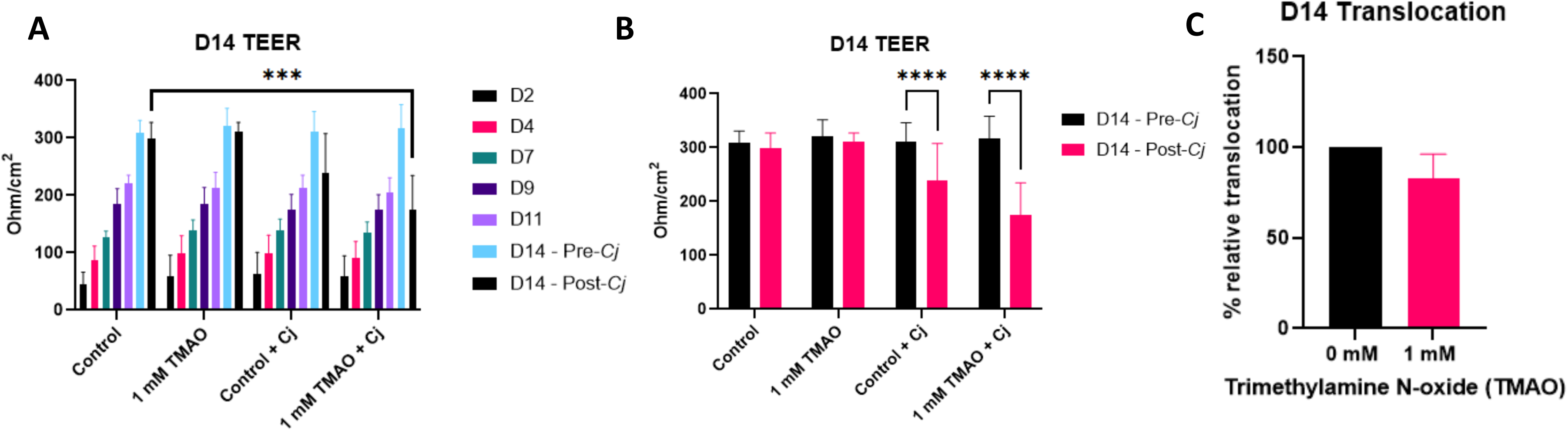
*C. jejuni* has significantly increased intestinal barrier damage in 14-day-old polarized Caco-2 cells. (A) The measured transepithelial electrical resistance (TEER) across the polarized epithelial layer over 14 days. (B) The measured TEER before and after microaerobic infection with *C. jejuni*. (C) The relative transmigration of *C. jejuni* 81-176 across the polarized monolayer after a 3-hour microaerobic infection. Data are presented as the mean ± SEM of three independent biological replicates. Statistical significance was assessed via (A) Two-Way RM ANOVA followed by Dunnett’s post hoc test, (B) Two-Way RM ANOVA followed by Sidak’s post hoc test, and (C) Student’s t-test. ** p<0.01, **** p<0.0001

**Supplemental Figure 5.**
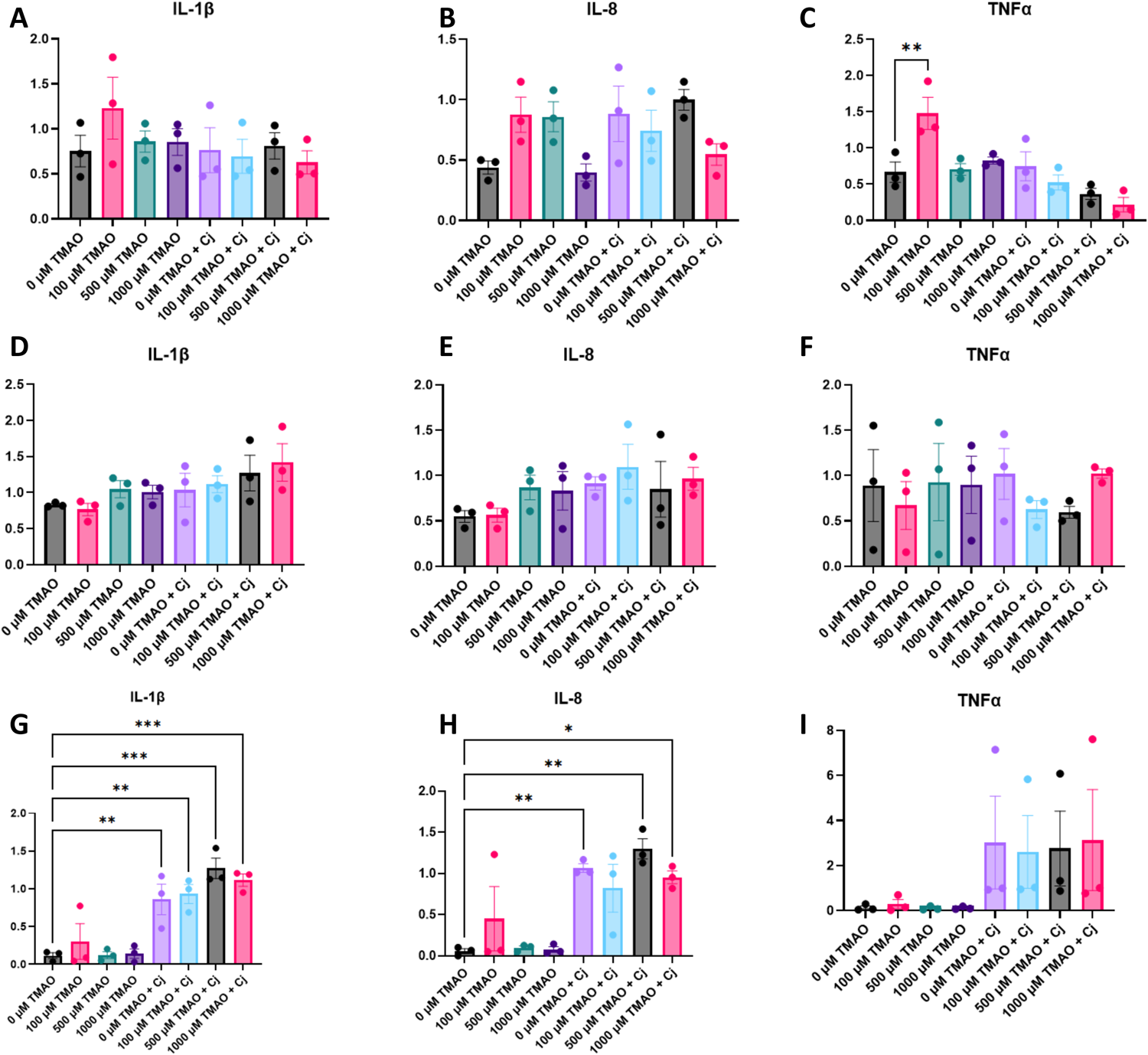
The inflammatory response of TMAO treated Caco-2 cell monolayers to *C. jejuni* after 3, 4, and 6 hours of infection. RT-qPCR of (A, D, G) IL-1β, (B, E, H) IL-8 and (C, F, I) TNFα of Caco-2 cells treated with TMAO for 48 hours at varied concentrations, and infected with *C. jejuni*. These infection assays were conducted over (A-C) 3 hours, (D-F) 4 hours, and (G-I) 6 hours. Monolayers were treated with TMAO and had either a terminal infection with *C. jejuni* for (A-C) 3 hours or (D-I) 4 hours under microaerobic conditions or incubated under microaerobic conditions for the same period of time. In panels (G-I), following their 4-hour infection, media in all wells were replaced with fresh media supplemented with 20 ug/mL gentamicin for 2 hours, under 5% CO_2_. Data are presented as the mean ± SEM of three independent biological replicates. Statistical significance was assessed via One-Way ANOVA followed by Sidak’s post hoc test.

